# Single-cell lineage tracing maps clonal and transcriptional dynamics in melanoma metastasis

**DOI:** 10.64898/2026.04.09.717571

**Authors:** Haiyin Li, Yeqing Chen, Jessica Kaster, Maggie Dunne, Cong Qi, Ling Li, Min Xiao, Monzy Thomas, Nazifa Promi, Dylan Fingerman, Gregory Schuyler Brown, Qiuxian Zheng, Jessie Villanueva, Bin Tian, Xiaowei Xu, Dave SB Hoon, Arjun Raj, Zhi Wei, Noam Auslander, Meenhard Herlyn

## Abstract

Melanoma metastasis is driven by extensive intratumoral heterogeneity and phenotypic plasticity, yet how clonal identity relates to transcriptional programs during metastasis remains unclear. Here, we applied MeRLin, a single-cell lineage tracing platform, to dissect the clonal and transcriptional heterogeneity of metastatic melanoma in a patient-derived spontaneous metastasis model. Clonal analyses revealed hierarchical structures during tumor progression, with a subset of lineages from primary tumors consistently enriched across metastatic sites, supporting a model of polyclonal seeding followed by selective expansion of pre-existing highly metastatic subpopulations. Single-cell transcriptomic profiling identified two major metastatic subpopulations of distinct transcriptional programs, characterized by neural crest stem cell-like and lipid metabolism signatures. Both programs were enriched for invasion-associated genes and maintained across organs through distinct regulatory networks. Spatial mapping by barcode RNA-FISH linked these transcriptional states to their tissue context and showed that *OLFML3* expression partially co-localized with a dominant subpopulation at the tumor-liver interface, marking the invasive fronts of metastatic growth. Together, these findings establish a framework in which clonal identity, transcriptional state, and spatial organization jointly shape metastatic melanoma progression.

## Introduction

Metastasis remains the major cause of melanoma-related mortality, in part because tumor dissemination is shaped by profound intratumoral heterogeneity and dynamic phenotypic plasticity^1^. Metastatic cells often exhibit organ-specific preferences, known as organotropism, consistent with the classical “seed and soil” hypothesis in which successful spread of cancer cells depends on both their intrinsic properties and the distal colonized microenvironment^2^. In melanoma, a key contributor to heterogeneity is phenotype switching, whereby tumor cells transition between more melanocytic states and more invasive, dedifferentiated states. Early gene expression profiling studies identified proliferative and invasive melanoma transcriptional states and demonstrated that melanoma cells can switch between these phenotypes *in vivo*, highlighting substantial transcriptional plasticity influenced by the tumor microenvironment^3^. Subsequent work revealed that melanoma cell state transitions are governed by distinct regulatory networks, including *SOX10/MITF*-driven proliferative programs and AP-1/TEAD-dependent invasive programs associated with metastasis and therapy resistance^4^.

Consistent with this model, single-cell RNA sequencing has revealed extensive intratumoral transcriptional heterogeneity in melanoma tumors, where malignant cells coexist in distinct transcriptional states characterized by high *MITF* expression or low *MITF* with elevated *AXL*, reflecting differences in cell-cycle activity, spatial context, and drug-resistance programs^5^. More broadly, multi-omic profiling of intrapatient metastatic melanoma has revealed a diverse evolutionary landscape. This includes polyclonal seeding and the distinct trajectory of brain metastases, which often diverge early in molecular evolution despite emerging clinically in later stages of disease^6^. Recent evidence further suggest that melanoma follows a hierarchical organization, wherein distinct cellular subpopulations contribute differentially to primary tumor expansion versus metastatic dissemination^7^. The identification of dedifferentiated neural crest-like subpopulations is particularly interesting, as it indicates that melanoma reactivates embryonic migration programs to drive cellular plasticity and invasion^8^. Despite these advances, how the interplay between clonal identity and microenvironmental adaptation contributes to phenotypic plasticity during metastatic dissemination remains incompletely understood.

To address this challenge, we applied MeRLin^9^, a barcoding-based single-cell lineage tracing platform, to dissect clonal dynamics and transcriptional states in metastatic melanoma using a patient-derived spontaneous metastasis model. By integrating lineage tracing with single-cell RNA sequencing and RNA fluorescence *in situ* hybridization, this approach enables simultaneous tracking of clonal fate, transcriptional programs, and tumor-microenvironment interactions across primary tumor and metastatic sites. We used this system to determine the prevalence of multi-organ disseminating clones and uncovered the phenotypic consistency of these highly metastatic subpopulations, characterized by neural crest stem cell-like (NC-like) and lipid metabolism-associated signatures. In addition, we identified a dominant metastatic subpopulation partially co-localized with a NC-like marker *OLFML3* at the tumor-liver interface *in situ*. Together, these findings provide new insights into how clonal identity, transcriptional state, and spatial organization interact to govern metastatic melanoma progression.

## Results

### Clonal dynamics of multi-organ melanoma dissemination

We applied the MeRLin high-complexity lentiviral barcoding system to the patient-derived melanoma cell line WND238 to trace the clonal dynamics of metastatic cells in a mouse model (Fig. 1A). The vector encodes firefly luciferase and mNeptune2.5, a far-red fluorescent protein optimized for deep-tissue imaging with minimal autofluorescence^9^ (Methods). Lineage tracing was performed by sequencing clone-specific 265 bp semi-random barcodes embedded within the 3′-untranslated region (UTR) of luciferase and mNeptune2.5 transcripts, allowing integrated analysis of clonal identity and transcriptional profiles at single-cell resolution^9^ (Fig. 1A; Methods). These transcribed barcodes also enabled targeted visualization of selected tumor subpopulations using RNA fluorescence *in situ* hybridization (RNA-FISH).

**Fig. 1.**
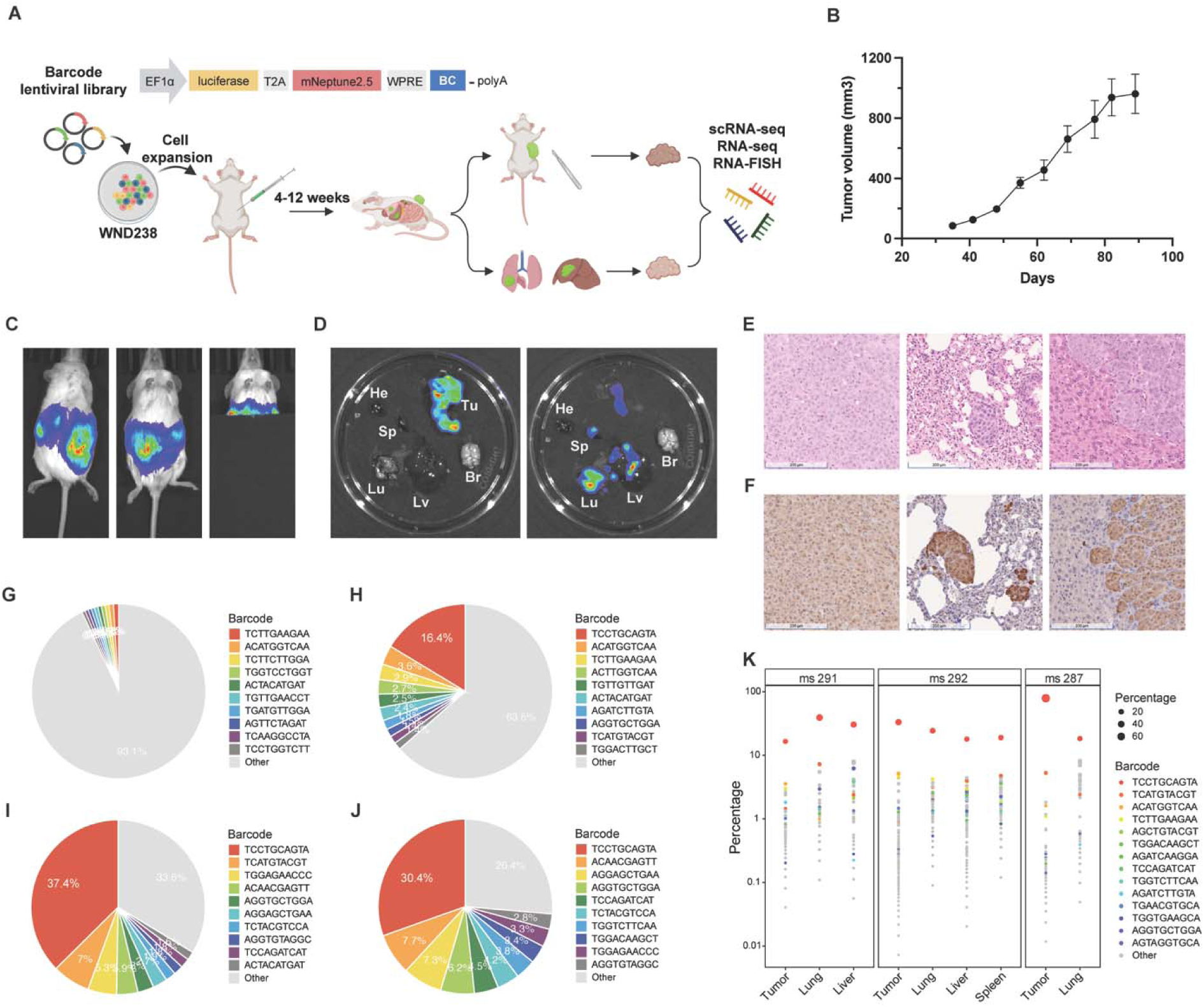
Lineage tracing of metastatic melanoma using the MeRLin barcoding system. **A**, Experimental outline. WND238 melanoma cells were transduced with the MeRLin lentiviral barcode library containing semi-random barcodes (BC) embedded within EF1α-driven luciferase and mNeptune2.5 transcripts. Barcoded cells were expanded *in vitro* and injected subcutaneously into NSG mice to establish tumors. Tumor growth and dissemination were monitored for 4 to 12 weeks. Primary tumors and metastatic lesions, including those from lung and liver, were excised for downstream analyses, including single-cell RNA sequencing (scRNA-seq), bulk RNA sequencing (RNA-seq), and RNA fluorescence in situ hybridization (RNA-FISH). **B**, *In vivo* growth curves of barcoded WND238 tumors (n = 3). **C**, Bioluminescence imaging (BLI) of a mouse bearing a barcoded WND238 tumor 83 days post injection. Signals are detected at the primary site in dorsal (left) and ventral (middle) views. After shielding the primary tumor with a light proof cover, residual signals are detected in the lungs (right). **D**, *Ex vivo* BLI of the excised primary tumor (Tu) and organs including lung (Lu), liver (Lv), spleen (Sp), brain (Br), and heart (He), imaged with (left) and without (right) the primary tumor. **E**, Hematoxylin and eosin (H&E) staining of sections from the primary tumor (left), lung (middle), and liver (right) of the second mouse (scale bar 200 μm). **F**, Immunohistochemical (IHC) staining for firefly luciferase in sections from the primary tumor (left), lung (middle), and liver (right) of the second mouse (scale bar 200 μm). **G**, Pie chart showing the clonal composition of the original control (Table S3, bulk RNA-seq). **H**, Pie charts indicating the hierarchical clonal composition of the primary tumor, lung (**I**), and liver (**J**) (Table S3, scRNA-seq). **K**, Bubble plot showing the clonal frequency of 14 barcodes shared across all primary and metastatic sites in three biological replicates (ms 291, ms 292, and ms 287).

WND238 was derived from a metastatic lesion of a 42-year-old male patient harboring a *BRAF* V600E mutation prior to therapy^10^. This model forms metastatic tumors spontaneously in immunodeficient mice under standard xenograft conditions (Table S1). WND238 cells were transduced with lentiviral barcode library at a multiplicity of infection (MOI) below 0.4 to ensure each cell carried a unique barcode (Fig. 1A). Transduced cells were sorted for mNeptune2.5-positive populations using flow cytometry. Barcoded cells were expanded and divided into biological replicates, with one replicate preserved as the original control. Replicate populations of MeRLin-barcoded WND238 cells were then injected subcutaneously into the right flanks of NSG (NOD-*scid* IL2Rgamma^null^) mice for tumor establishment (Methods).

Twelve weeks post injection, primary tumors reached ∼1,000 mm³ (Fig. 1B). *In vivo* bioluminescence imaging (BLI) detected signals at the primary sites, with mice imaged in both dorsal and ventral views (Fig. 1C left and middle, Fig. S1C, and S1F). Shielding of the primary tumor revealed thoracic signals consistent with lung metastases. (Fig. 1C right). In parallel, *in vivo* fluorescence imaging (ex/em 640/680 nm) confirmed signals from primary tumors, but with a lower signal-to-noise ratio than BLI (Fig. S1A, S1C, and S1F).

Mice were euthanized and primary tumors and organs (lung, liver, spleen, brain, and heart) were excised for analysis. *Ex vivo* BLI detected signals in the primary tumor, lung, liver, and spleen (Fig. 1D). A second mouse showed a similar distribution with the liver exhibiting the highest metastatic signal (Fig. S1D), whereas the third mouse displayed predominantly lung-localized signals (Fig. S1G). Far-red fluorescence imaging confirmed these patterns, albeit with lower resolution and sensitivity than bioluminescence (Fig. S1B, S1E, and S1H).

Hematoxylin and eosin (H&E) staining of primary tumor, lung, and liver sections confirmed histological features of metastatic lesions (Fig. 1E). In the second mouse, liver metastases formed invasive fronts of densely packed malignant cells infiltrating liver parenchyma (Fig. 1E). Immunohistochemical (IHC) staining for firefly luciferase further verified the presence of MeRLin-barcoded cells across these sites (Fig. 1F).

Mouse tissues were dissociated and enriched for viable melanoma cells by depleting murine and red blood cells (Methods). Single-cell RNA sequencing (scRNA-seq) was performed on the primary tumor, lung, and liver from the first mouse, while bulk RNA-seq was conducted on the original control and all tumor and metastatic sites from the remaining mice (Table S2). After quality control filtering to remove duplicates and artifacts, barcode reads extracted from scRNA-seq were used to quantify clonal composition across sites in the first mouse, whereas barcode reads from bulk RNA-seq were used to estimate clonal composition in the original control and samples from the other two mice (Table S3; Methods). In the WND238 metastatic model, barcode distributions became progressively skewed during tumor progression. While the original control showed a relatively even distribution (Fig. 1G), the primary tumor exhibited a hierarchical clonal composition (Fig. 1H). This contrasts with the non-metastatic WM4237-1 model^9^, which retained comparatively even distributions in both control cells and engrafted tumors (Fig. S2B-C). This hierarchical shift was more pronounced in WND238 lung and liver metastases, where the ten most abundant lineages accounted for 66.4% and 73.6% of detected barcodes, respectively (Fig. 1I-J).

Bulk RNA-seq analyses of the remaining two mice showed similarly skewed, hierarchical clonal compositions across primary tumors and metastatic sites (Fig. S2D-I). Notably, fourteen barcodes were shared across all sites in the three biological replicates (Fig. 1K), indicating that these lineages consistently contributed to both tumor growth and dissemination. Together, these results demonstrate substantial clonal heterogeneity alongside reproducible enrichment of a subset of dominant lineages, consistent with polyclonal seeding despite inter-mouse variability.

### Single-cell lineage tracing reveals clonal heterogeneity

Uniform manifold approximation and projection (UMAP) of single-cell transcriptomes revealed distinct transcriptional states of melanoma cells within the primary tumor, lung, and liver metastases (Fig. 2A). Cells from the primary tumor formed two central clusters, from which metastatic populations extended into two lung-enriched transcriptional clusters and further into adjacent liver-enriched clusters, suggesting progressive transcriptional divergence associated with metastatic dissemination and colonization. We matched barcode sequences identified in lung and liver metastases to those present in the primary tumor (Fig. 2B). Among 403 barcode-defined subpopulations detected in the primary tumor, 34 clones were found exclusively in the lung, 20 exclusively in the liver, and 45 in both organs, while the remaining 304 subpopulations were detected only in the primary tumor, enabling classification of subpopulations based on their metastatic fates.

**Fig. 2.**
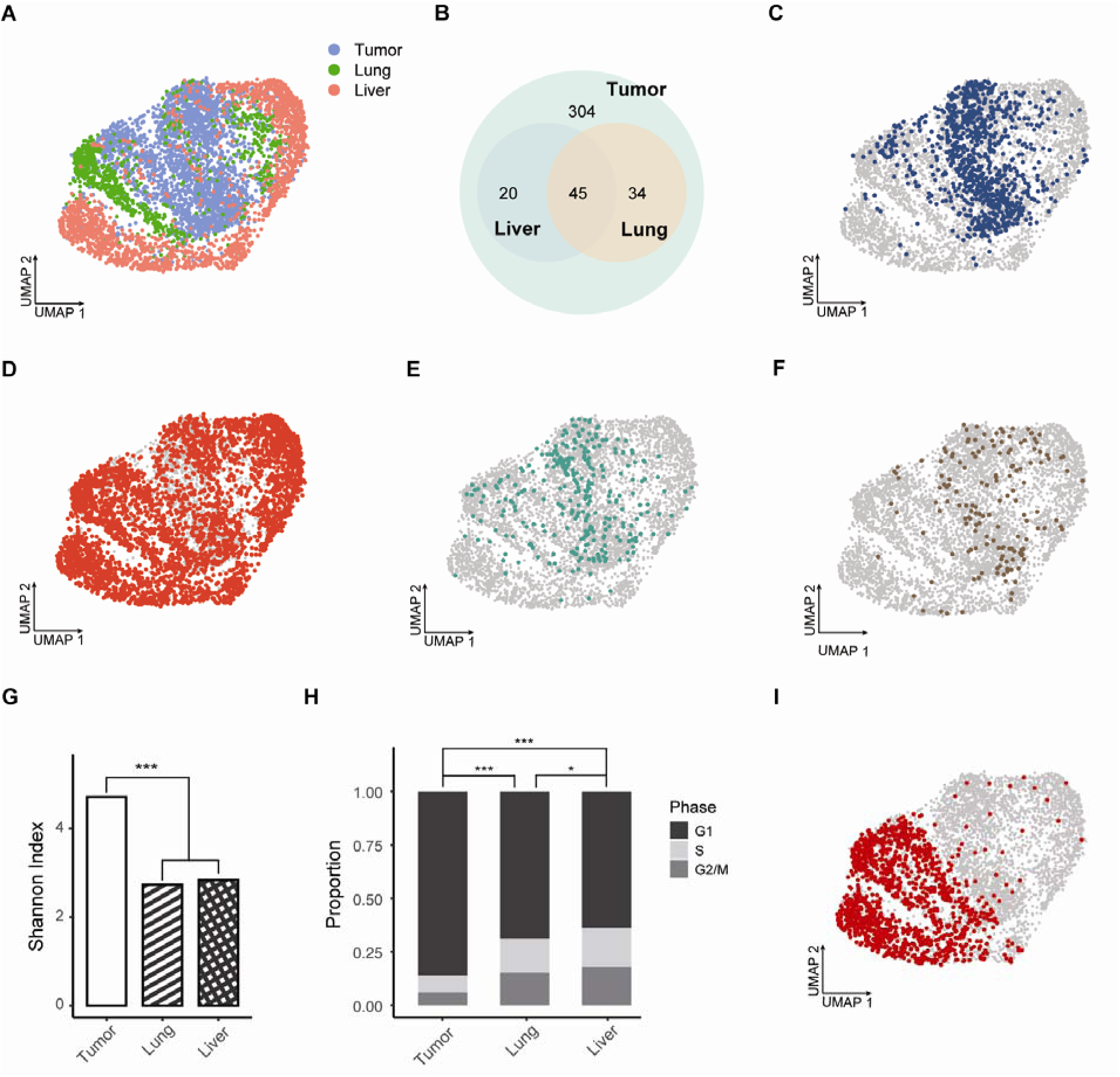
Clonal dissemination from the primary tumor to lung and liver metastases. **A**, UMAP visualization of single-cell transcriptomes of melanoma cells from the primary tumor (blue), lung (green), and liver (red). **B**, Venn diagram showing unique barcodes identified by scRNA-seq in the primary tumor, lung, and liver metastases. **C**, UMAP highlighting 304 non-metastatic subpopulations restricted to the primary tumor. **D**, UMAP showing 45 shared metastatic subpopulations shared between lung and liver. **E**, UMAP showing 34 metastatic subpopulations exclusive to lung. **F**, UMAP showing 20 metastatic subpopulations exclusive to liver. **G**, Shannon diversity index comparing barcode diversity of cells from the primary tumor, lung, and liver metastases. *** *P* < 0.001, two-tailed Hutcheson’s *t*-test. **H**, Distribution of cell cycle phases inferred from scRNA-seq. * *P* < 0.05, *** *P* < 0.001, two-tailed Fisher’s exact test. **I**, UMAP visualization showing the metastatic fate of the most abundant subpopulation across sites.

The 304 tumor-only subpopulations were predominantly localized within one of the two central clusters (Fig. 2C), indicating that this transcriptional state is largely associated with non-metastatic behavior. In contrast, the 45 highly metastatic subpopulations were distributed across all clusters and partially overlapped with the tumor-only populations (Fig. 2D), consistent with increased transcriptional heterogeneity among clones capable of disseminating to both organs. Lung-only and liver-only metastatic subpopulations were mainly derived from one of the central clusters (Fig. 2E-F), which substantially overlapped with tumor-only subpopulations (Fig. 2C), suggesting a closer transcriptional relationship between these groups. Shared metastatic subpopulations predominated, accounting for over 80% of cells in lung metastases and over 90% in liver metastases (Fig. S3A).

The number of unique barcodes detected in lung and liver metastases was comparable (Fig. S3B), while the total number of barcodes recovered from the liver was approximately two-fold higher than that from the lung (Fig. S3C). Shannon index which quantifies barcode richness and evenness^11^ (Methods), showed a marked and comparable reduction in barcode diversity in both metastatic sites relative to the primary tumor (Fig. 2G), reflecting substantial clonal restriction associated with metastatic dissemination and outgrowth. Cell cycle phase inferred from scRNA-seq data revealed a significantly higher proportion of cycling cells in the G2/M and S phases in both lung and liver metastases (Fig. 2H; Methods), with liver metastases exhibiting the greatest proliferative fraction, while cells in the primary tumor exhibited G1 cell cycle arrest.

The most abundant subpopulation in the primary tumor, representing 16.4% of all cells (Fig. 1H), expanded to 37.4% and 30.4% in lung and liver metastases, respectively (Fig. 1I–J), indicating preferential representation of this clone in both organs. This subpopulation was enriched in one of the two central clusters and extended broadly into one of the two lung-enriched clusters and its adjacent liver-enriched cluster (Fig. 2I). In contrast, the subpopulations ranked 2 to 5 in the primary tumor was preferentially localized to the other central cluster (Fig. S3D), suggesting that dominant clones occupying distinct transcriptional states contribute to metastatic heterogeneity.

### Clonal and transcriptional dynamics of melanoma metastasis

We analyzed alternative polyadenylation (APA) isoform expression from scRNA-seq data, a widespread regulatory mechanism that generates distinct 3′ transcript ends and is closely linked to cell identity, proliferation, and differentiation^12^ (Methods). Comparative analyses revealed increased frequencies of both 3′UTR lengthening and shortening events in lung and liver metastases as compared to the primary tumor, with liver metastases exhibiting a predominant global shift toward shorter 3′UTRs (Fig. 3A). These results suggest dynamic post-transcriptional regulation during melanoma dissemination to distant organs.

**Fig. 3.**
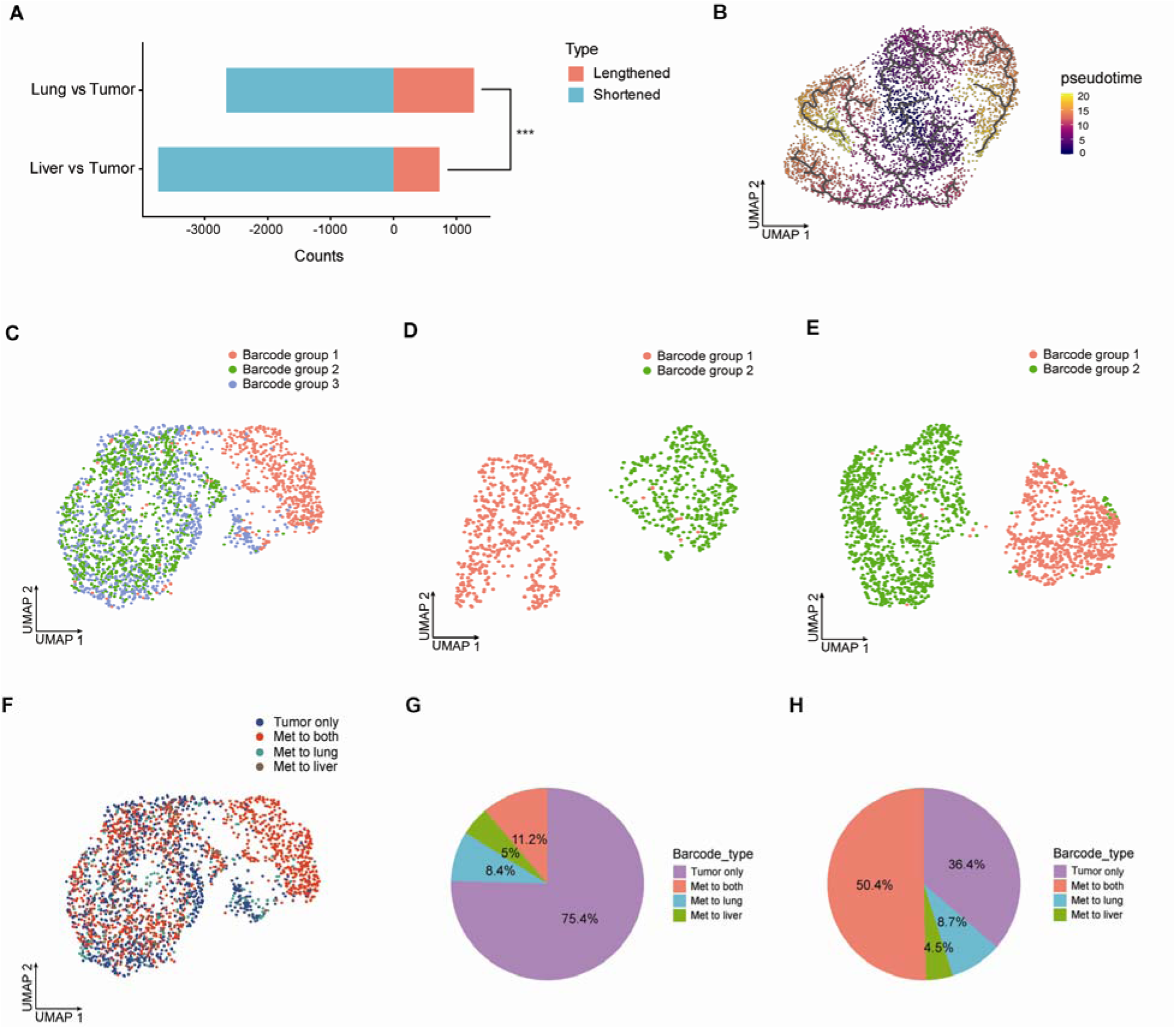
Clonal and transcriptional dynamics of melanoma metastasis. **A**, Alternative polyadenylation (APA) analysis showing changes in 3′UTR length distributions in lung and liver metastases compared with the primary tumor. *** *P* < 0.001; two-tailed Fisher’s exact test. **B**, UMAP projection of single cells colored by inferred pseudotime values, with black curves indicating the inferred trajectory backbone. **C**, UMAP visualization of hybrid clusters integrating clonal and transcriptomic information from scRNA seq in the primary tumor, lung metastases (**D**), and liver metastases (**E**). **F**, UMAP showing barcode types classified by metastatic fate in the primary tumor. **G**, Pie chart showing the barcode types by their metastatic fates in the primary tumor. **H**, Pie chart showing cell numbers of barcode types classified by metastatic fate in the primary tumor.

To characterize transcriptional relationships associated with metastatic progression, we performed unsupervised pseudotime analysis (Methods), which revealed that metastatic cells were organized along two major transcriptional trajectories connected to the primary tumor population (Fig. 3B). The first trajectory overlapped extensively with one of the central clusters and extended into a continuous, branched structure giving rise to lung- and liver-enriched branches. In contrast, the second trajectory segregated into lung- and liver-enriched branches positioned on opposing arms. Together, these results indicated that metastatic cells follow diverse transcriptional trajectories relative to the primary tumor.

We applied ClonoCluster^13^, a clustering algorithm that integrates clonal identity and transcriptomic similarity, to generate hybrid clusters from scRNA-seq data independently at each tumor site, thereby improving cluster resolution and facilitating marker identification by adjusting the contribution of clonal barcode identity within the UMAP embedding^9^ (Fig. S4A-C; Methods). In the primary tumor, ClonoCluster resolved three major barcode-associated groups (Fig. 3C), with group 1 showing clear separation from the remaining groups, while groups 2 and 3 exhibited substantial overlap in transcriptional space. In lung and liver metastases, two barcode-associated groups were identified in each organ (Fig. 3D-E).

The most abundant barcode suffix “TCCTGCAGTA” in the primary tumor (Fig. 1H) was also the dominant constituent of group 1 across both organs (Fig. S4D), indicating a strong correspondence between clonal abundance in the primary tumor and representation at metastatic sites. Similarly, the 14 shared barcodes (Fig. 1K) accounted for a substantial fraction of barcode reads across primary and metastatic samples (Fig. S4E). Together, these observations indicate that although multiple clonal lineages contribute to metastatic dissemination, a subset of highly represented clones is consistently detected across metastatic sites.

Mapping metastatic fate onto the UMAP of primary tumor showed that cells contributing to both lung and liver metastases were predominantly localized within groups 1 and 2 (Fig. 3F). Quantitatively, lineages shared between lung and liver metastases comprised 11.2% of the clones within primary tumor, while lung-only and liver-only metastatic lineages accounted for 8.4% and 5%, respectively (Fig. 3G). In terms of cell numbers, metastatic subpopulations shared between lung and liver accounted for 50.4% of primary tumor cells, while lung-only and liver-only subpopulations comprised 8.7% and 4.5%, respectively. In contrast, non-metastatic subpopulations, despite representing 75.4% of clonal lineages, accounted for only 36.4% of primary tumor cells (Fig. 3H).

### Selected genes govern melanoma metastasis

To identify the cellular expression programs associated with metastatic subpopulations, we performed differential gene expression (DGE) analysis for each barcode group independently at each tumor site (Table S4; Methods). Comparative analysis between barcode groups 1 and 2 revealed distinct yet highly consistent transcriptional programs across primary and metastatic sites. In total, 91 genes upregulated in barcode group 1 were shared across all three sites, while 64 genes upregulated in barcode group 2 were similarly conserved across sites (Table S4), indicating that these transcriptional states are largely maintained during metastatic dissemination.

Among the shared genes, the melanoma dedifferentiation marker *AXL* emerged as one of the most significantly enriched transcripts in barcode group 1 across primary tumor and metastatic lesions^14^ (Fig. 4A; Table S4). This enrichment is consistent with barcode group 1 cells adopting a relatively invasive and dedifferentiated phenotype that persists as cells disseminate to distant organs. Concordantly, multiple established melanoma invasion and metastasis genes, including *NEDD9*^15^, *SERPINA3*^16,17^, *TRPV4*^18^, *NOX4*^19^, *PXN*^20^, and *EFNA5*^21^, were significantly enriched in barcode group 1 across sites (Fig. 4C and Fig. S5C-D). In addition, invasion-associated genes such as *ZEB1*^22,23^, *CDCP1*^24^, *CDC42EP5*^25^, *BACH1*^26^, and *IGF2BP3*^27^ were selectively upregulated in barcode group 1 within lung and liver metastases (Fig. 4D and 4C), suggesting further reinforcement of invasive programs at secondary sites.

**Fig. 4.**
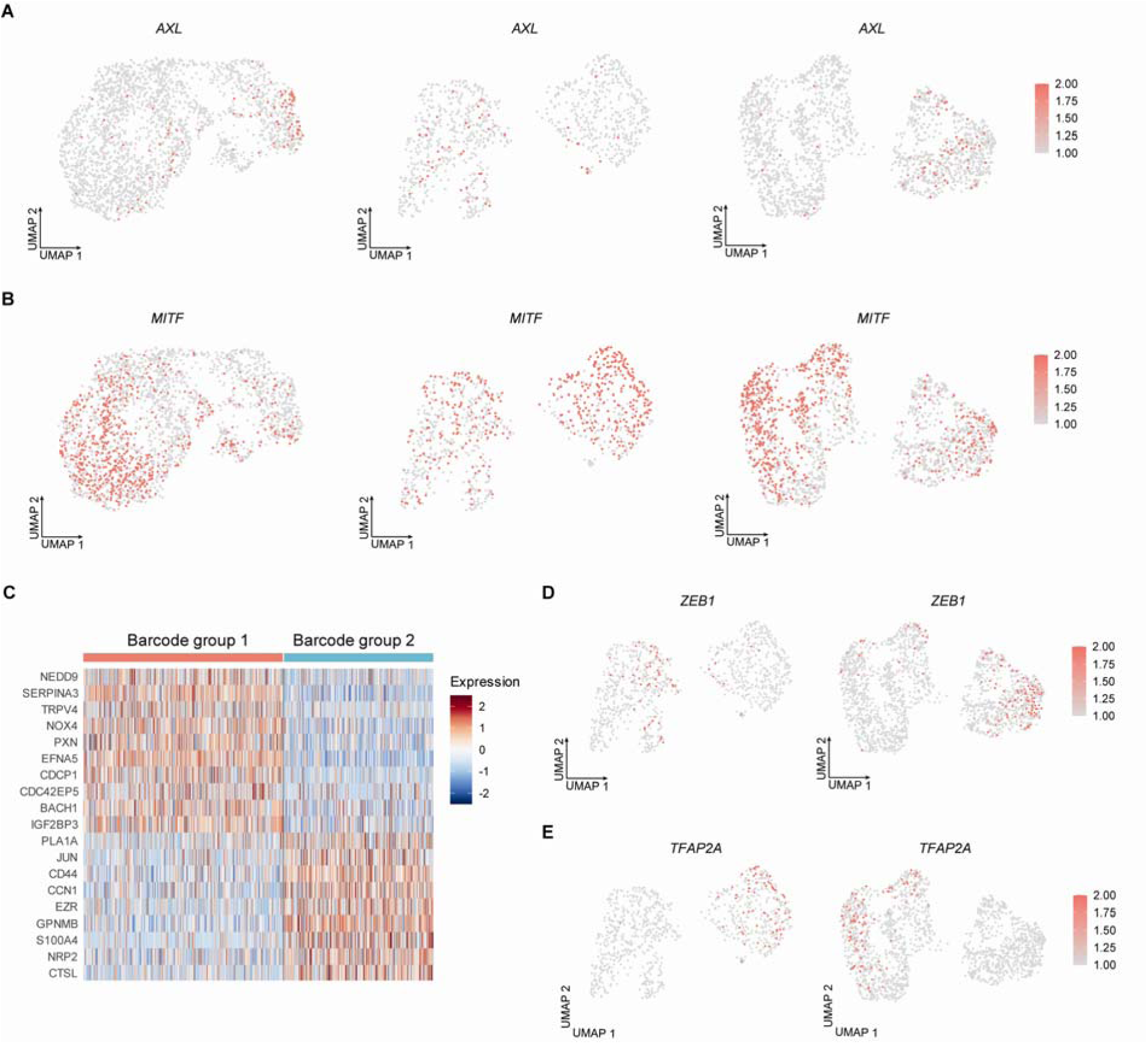
Expression of dedifferentiation, melanocytic, and invasion-associated genes in metastatic melanoma. **A**, UMAP visualization of expression of the melanoma dedifferentiation marker *AXL* in the primary tumor (left), lung metastases (middle), and liver metastases (right). **B**, UMAP visualization of expression of the melanocytic marker *MITF* in the primary tumor (left), lung metastases (middle), and liver metastases (right). **C**, Heatmap showing melanoma invasion-associated genes identified for each barcode group in lung metastases (Table S4). Fold change ≥ 1.5, cell percentage ≥ 40%, and adjusted *P* < 0.05. **D**, UMAP visualization of *ZEB1* expression in lung metastases (left) and liver metastases (right). **E**, UMAP visualization of *TFAP2A* expression in lung metastases (left) and liver metastases (right).

In contrast, *MITF*, a master regulator of melanoma differentiation^28^, was significantly enriched in barcode group 2 cells across the primary tumor as well as lung and liver metastases (Fig. 4B; Table S4). This pattern indicates that, relative to barcode group 1, these metastatic subpopulations exist in a more differentiated melanocytic state and maintain this identity during dissemination. Consistently, melanocytic marker *PMEL* was enriched in barcode group 2 across tumor sites (Fig. S5A), while another differentiation marker *DCT* was significantly upregulated in barcode group 2 specifically in lung and liver metastases but not in primary tumor (Fig. S5B; Table S4).

Notably, despite their differentiated transcriptional profile, barcode group 2 cells also exhibited enrichment of genes of melanoma invasion and metastasis, including *PLA1A*^29^, *JUN*^30^, *CD44*^31^, *CCN1*^32^, *EZR*^33^, and *GPNMB*^34^, across all sites (Fig. 4C and and Fig. S5C-D). Moreover, invasion-associated genes such as *S100A4*^35^, *NRP2*^36^, *CTSL*^37^, *MARCKS*^38^, and *TFAP2A*^39^ were selectively enriched in barcode group 2 within lung and liver metastases (Fig. 4E and 4C). Together, these findings suggest that metastatic subpopulations can sustain differentiated melanocytic programs while concurrently activating invasion and metastasis-associated pathways, highlighting the coexistence of phenotypic differentiation and invasive potential.

### NC-like and lipid metabolism programs define metastatic subpopulations

EnrichR pathway analysis was performed to functionally annotate each hybrid cluster (Fig. 5A; Table S5). Hallmark, KEGG, Reactome, and Wikipathway databases were queried to identify biological processes associated with each group’s transcriptional program (Table S6; Methods). The activities of these expression programs were quantified using AUCell and projected onto UMAP space (Methods). This analysis showed that group 1 was enriched for neural crest stem-like (NC-like) features^40^ (Fig. 5B-D top), while group 2 was preferentially associated with lipid metabolism-related pathways^41^ (Fig. 5B-D bottom). Notably, the most abundant barcode aligned with the NC-like signature across all sites (Fig. S4D).

**Fig. 5.**
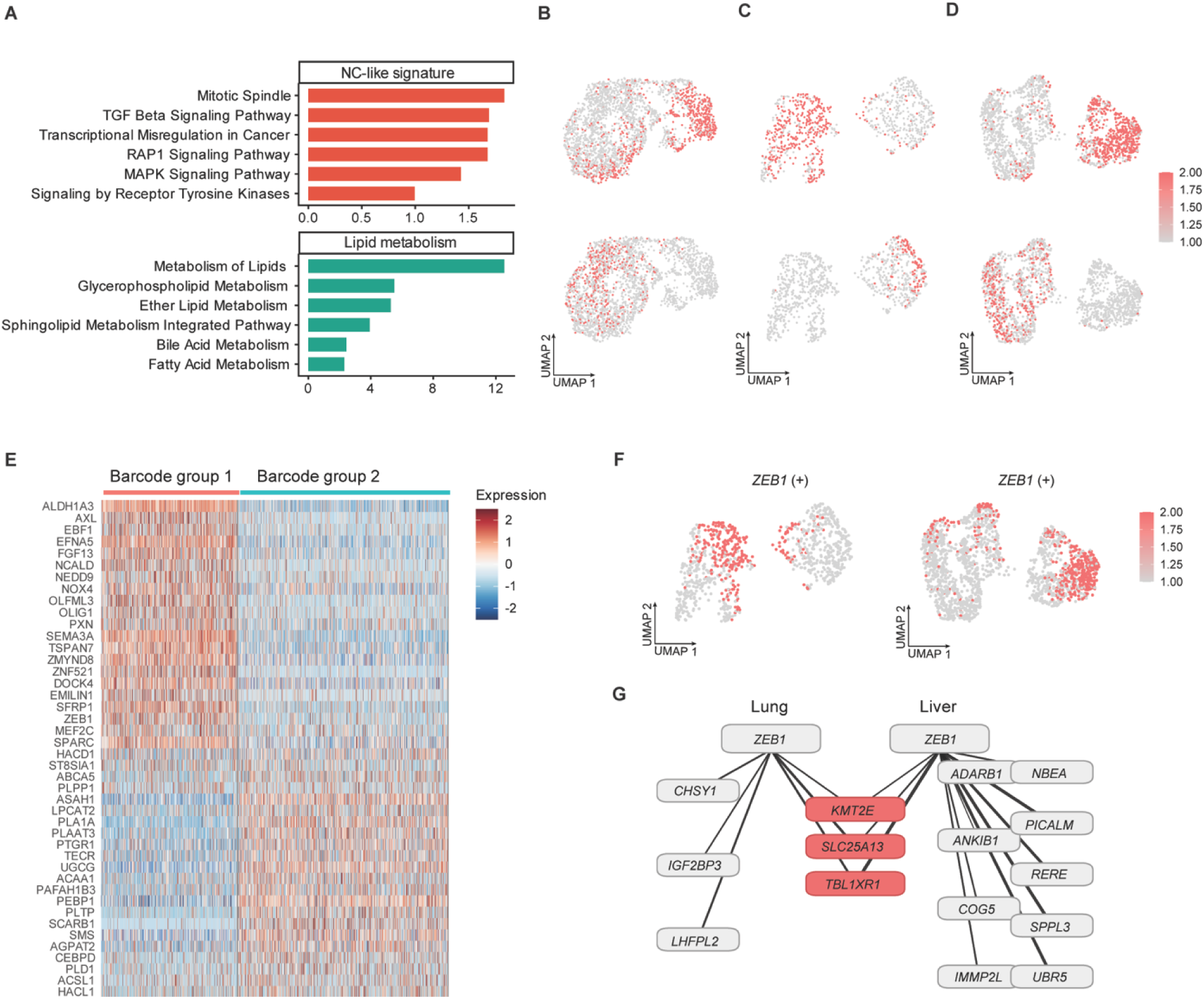
Distinct transcriptional programs define metastatic subpopulations. **A**, Functional enrichment terms for barcode groups 1 and 2 identified in Fig. 3C to 3E. *P*-values were determined using two-tailed Fisher’s exact tests (Table S6). **B**, AUCell scores (color scale) for enriched functional programs in each barcode group projected onto UMAP from the primary tumor, lung metastases (**C**), and liver metastases (**D**). Top, neural crest stem-like (NC-like) signature; bottom lipid metabolism signature. **E**, Discriminative marker genes identified for each barcode group in liver metastases (Table S5). Fold change ≥ 1.5, cell percentage ≥ 40%, and adjusted *P* < 0.05. **F**, SCENIC analysis showing regulon activity of transcription factor *ZEB1* projected onto the UMAPs from lung metastases (left) and liver metastases (right). **G**, Predicted target genes of *ZEB1* in the NC-like subpopulations of lung and liver metastases.

The metastatic cells from barcode group 1 exhibited an NC-like transcriptional program characterized by increased expression of *NEDD9*, *ALDH1A3*, *EFNA5*, *SEMA3A*, *OLFML3*, *PXN*, and *NOX4* across primary and metastatic sites (Fig. 5E and Fig. S6A-B). In addition, elevated expression of *EMILIN1*, *DOCK4*, and *SFRP1* was observed in barcode group 1 cells within lung and liver metastases (Table S5), while *SPARC* and *MEF2C* showed preferential upregulation in barcode group 1 cells in liver metastases (Fig. 5E). In contrast, cells from barcode group 2 displayed enrichment of lipid metabolism-associated genes, including *ASAH1*, *LPCAT2*, *UGCG*, *PLAAT3*, and *TECR*, across all sites (Fig. 5E and Fig. S6A-B). Additional lipid metabolism-related genes in barcode group 2 exhibited site-associated expression patterns, with *ACAA1*, *PLTP*, *SCARB1*, and *SMS* elevated in both lung and liver metastases (Table S4), *ACSL1* and *HACL1* preferentially increased in lung metastases (Fig. S6B), and *AGPAT2* and *PLD1* selectively increased in liver metastases (Fig. 5E).

Single-cell regulatory network inference and clustering (SCENIC) analysis distinguished the regulatory mechanism associated with the NC-like and lipid metabolism programs across primary and metastatic sites (Fig. S7A-C; Methods). Regulon activities of *ZEB1*, *LEF1*, and *RB1* were consistently elevated in NC-like subpopulations within lung and liver metastases but were not detected in NC-like cells in the primary tumor (Fig. 5F and Fig. S7D-E). In contrast, *JUND* regulon activity was enriched in lipid metabolism-associated subpopulations across all sites (Fig. S7F), while *MITF* and *TFAP2A* activities were selectively increased in lung and liver metastases (Fig. S7G-H). Unsupervised analysis identified sets of *ZEB1* target genes in lung and liver metastases independently, with *KMT2E*, *SLC25A13*, and *TBL1XR1* shared between the two sites (Fig. 5G; Table S7). These results indicate site-specific *ZEB1* regulatory activity, with distinct subsets of target genes preferentially upregulated in lung versus liver metastases.

### Cell-cell communication and spatial mapping of metastatic subpopulations

CellChat analysis was performed to infer ligand-receptor-mediated communication between barcode-defined subpopulations across primary and metastatic sites (Fig. 6A; Methods). In the primary tumor, inferred signaling through SPP1, Midkine (MK), and collagen pathways showed predominant inter-group communication from barcode group 1 to barcode groups 2 and 3, together with detectable intra-group communication within barcode group 1. Relative lower SPP1 signaling was observed from barcode group 3 to barcode groups 1 and 2, and lower collagen-associated signaling was detected from barcode group 3 to barcode group 1. Laminin-mediated communication was comparatively weaker within barcode group 2 and in interactions from barcode group 2 to barcode group 3.

**Fig. 6.**
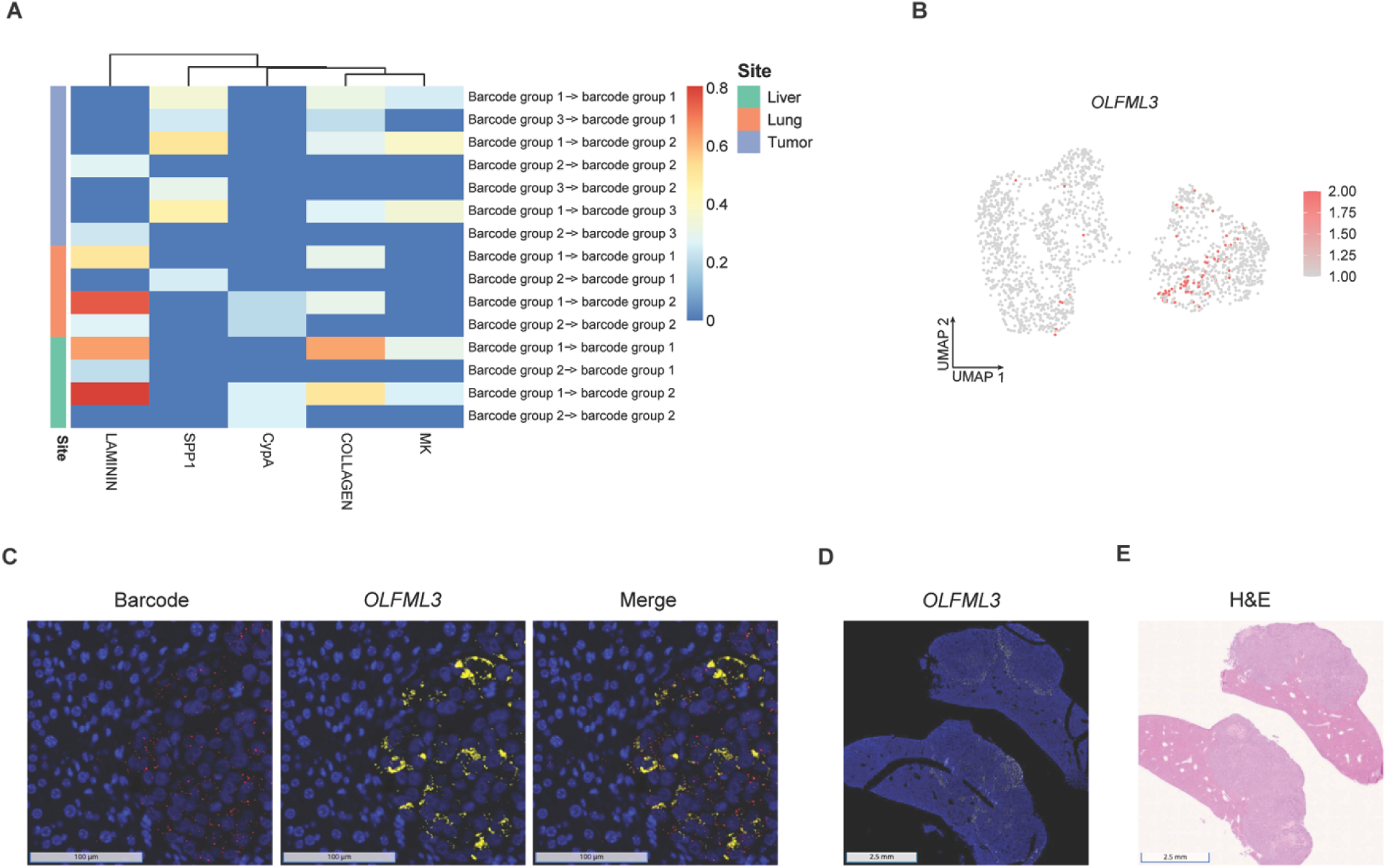
Cell-cell communications and spatial localization of metastatic subpopulations. **A**, Heatmap showing the relative contribution of SPP1, Midkine (MK), collagen, laminin, and CypA to outgoing (ligand to receptor) and incoming (receptor to ligand) interactions between barcode groups across the primary tumor (purple), lung (orange), and liver (green) metastases. **B,** UMAP projection showing expression of the NC-like marker *OLFML3* in liver metastases. **C,** Barcode RNA-FISH showing spatial localization of a dominant metastatic subpopulation carrying barcode suffix “TCCTGCAGTA” (red, left), *OLFML3* expression (yellow, middle) and merged partial co-localization (right) in liver metastases (scale bar 100 μm). DAPI staining was used to visualize cell nuclei (blue). **D,** RNA-FISH showing **OLFML3** expression (yellow) in liver tissue at low magnification (scale bar 2.5 mm). **E,** H&E staining of liver tissue sections (scale bar 2.5 mm).

In lung and liver metastases, inferred communication patterns were dominated by extracellular matrix-associated pathways, most notably laminin and collagen signaling. In lung metastases, laminin-associated signaling was enhanced, particularly in interactions with barcode group 1 as the sender and barcode group 2 as the receiver. In liver metastases, laminin signaling represented the most prominent inferred communication program, with directional interactions largely involving barcode group 1 as the sender. Collagen-mediated signaling was also increased in liver metastases, especially within intra-group communication of barcode group 1. MK-associated signaling remained detectable with barcode group 1 as the sender in liver metastases but showed relatively lower strength compared with the primary tumor. CypA-mediated signaling was present at low levels in both lung and liver metastases, while SPP1 signaling was more limited in metastatic sites compared to the primary tumor.

Across the primary and metastatic sites, barcode group 1 consistently exhibited stronger outgoing signaling in both inter-group and intra-group interactions compared with barcode groups 2 and 3. Together, these observations indicate that melanoma cell-cell communication in metastases is predominantly associated with extracellular matrix-related pathways, with site-dependent variation in their contributions during metastatic progression.

MeRLin-enabled barcode RNA-FISH provides an approach to assess clonal identity and gene expression *in situ*. We recovered the full-length barcode sequence of a selected subpopulation from bulk RNA-seq data (Methods). RNA-FISH probes were designed to target the barcode suffix “TCCTGCAGTA”, corresponding to the most abundant subpopulation detected across primary and metastatic sites (Fig. 1H-J). Based on scRNA-seq analyses, this barcode maps to the NC-like state in barcode group 1 (Fig. S4D). Analysis of scRNA-seq data showed that the NC-like marker gene *OLFML3* was significantly upregulated in barcode group 1 across the primary tumor, lung and liver metastases (Fig. S8A-B and 6B). Within liver metastases, *OLFML3* expression was enriched in a subset of barcode group 1 cells (Fig. 6B), which partially overlapped with cells carrying the barcode suffix “TCCTGCAGTA” (Fig. S4D right).

Using RNA-FISH (Methods), we spatially mapped this dominant metastatic subpopulation carrying barcode suffix “TCCTGCAGTA” together with *OLFML3* expression in liver metastases from the second mouse. Most melanoma cells at the invasive front showed positive staining for the barcode RNA-FISH probes (Fig. 6C, left), whereas non-specific negative control probes showed only weak background staining in mouse hepatocytes and did not label melanoma cells (Fig. S8C). *OLFML3* RNA-FISH signal was enriched at the invasive front (Fig. 6C, middle) and spatially overlapped with the barcode signal (Fig. 6C, right), while probes targeting human *AXL* produced weak background staining in mouse hepatocytes but no detectable signal in melanoma cells (Fig. S8D). Due to its relatively high expression, *OLFML3* RNA-FISH signal was detectable at low magnification (Fig. 6D). At this scale, *OLFML3* expression was localized along the tumor-liver interface, outlining the invasive fronts across tumor mass, consistent with H&E staining (Fig. 6E). Similar spatial patterns of the barcode and *OLFML3* expression were observed at invasive fronts in additional regions of interest (ROI) (Fig. S6E-F).

## Discussion

Most cancer-associated deaths result from metastasis, yet metastasis remains incompletely understood as an evolving, heterogeneous, and systemic disease. Its progression requires the stepwise acquisition of traits that enable dissemination, driven by clonal selection and the ability of metastatic cells to dynamically transition between distinct cellular states^42^. Here, we applied MeRLin^9^, a single-cell lineage tracing platform, to establish a framework for understanding how clonal identity, transcriptional state, and spatial organization collectively shape metastatic behavior in a patient-derived spontaneous melanoma metastasis model.

MeRLin lineage tracing revealed clonal barcode compositions that refine the classical “seed and soil” hypothesis. We observed a pronounced clonal hierarchy, in which a limited subset of lineages consistently contributed to primary and metastatic sites, indicating that metastatic competence is largely restricted to pre-existing “seeds” with high metastatic fitness. This pattern is consistent with a prior lineage reconstruction study in metastatic pancreatic cancer that demonstrated extensive bottlenecking, subpopulation signaling, and distinct transcriptional states associated with metastatic aggressiveness^43^. Moreover, the coexistence of shared and organ-specific subpopulations across lung and liver metastases supports a model of polyclonal dissemination shaped by selection from organ-specific microenvironments, consistent with previous observations of clear lineage segregation across melanoma metastases sampled at autopsy^44^.

Our data further suggest that metastatic outgrowth is governed by dynamic interactions between intrinsic transcriptional programs and tissue niches. Metastatic cells diverged from the primary tumor along distinct transcriptional trajectories, accompanied by site-specific alternative polyadenylation changes, indicating both transcriptional and post-transcriptional adaptation. In parallel, cell-cell communication shifted from diverse signaling in the primary tumor to extracellular matrix-associated programs in metastases, with enriched laminin and collagen signaling and the dominant clonal populations driving both inter-group and intra-group interactions. Together, these findings support a model in which intrinsic cellular states and extrinsic microenvironmental cues together shape metastatic progression.

Integrated single-cell analyses indicated that metastatic melanoma comprised co-existing cellular states that preserved lineage identity while acquiring invasive capacity, highlighting that differentiated melanocytic states and metastatic potential were not mutually exclusive. Distinct subpopulations maintained stable yet divergent transcriptional programs across sites, including an NC-like, invasion-associated state and a lipid metabolism-associated state, both of which were conserved during dissemination and further refined in a site-specific manner.

The NC-like program was characterized by upregulation of *AXL*, *NEDD9*, *ALDH1A3*, *EFNA5*, *SEMA3A*, *OLFML3*, *PXN*, and *NOX4*, with additional enrichment of *EMILIN1*, *DOCK4*, *SFRP1*, *SPARC*, and *MEF2C* in metastatic contexts^27^, whereas the lipid metabolism program, marked by *ASAH1*, *LPCAT2*, *UGCG*, *PLAAT3*, and *TECR*, was maintained across sites with organ-specific adaptations, including preferential upregulation of *ACSL1* and *HACL1* in lung metastases and *AGPAT2* and *PLD1* in liver metastases. These findings indicated functional specialization of metastatic lineages across tissue environments. Consistently, SCENIC analysis suggested that these states were governed by distinct yet partially conserved transcriptional networks, with *ZEB1*, *LEF1*, and *RB1* regulons enriched in NC-like subpopulations in metastases, indicating that this state was further remodeled in metastatic contexts rather than simply carried over from the primary tumor. In contrast, the lipid metabolism-associated population retained conserved *JUND* regulon activity across all sites, consistent with a more stable yet metastatic differentiated program. Together, these results support a model in which metastatic progression is driven by stable clonal compositions that dynamically engage invasion-associated pathways in a context-dependent manner.

Our results align with a cooperative model of melanoma metastasis in which proliferative and invasive cells form structured heterotypic clusters that cooperate in metastatic seeding, with *TFAP2A/E* acting as a master regulator of neural crest-like programs^45^. We observed that distinct clonal subpopulations coexisted within primary tumors and metastases, with NC-like populations exhibiting stronger outgoing signaling across sites. MeRLin-enabled barcode RNA-FISH further demonstrated that a dominant metastatic subpopulation associated with NC-like state co-expressed *OLFML3* at the invasive front of liver metastases. These findings suggest that invasive, dedifferentiated cells may actively remodel the tumor microenvironment to facilitate colonization, while more melanocytic subpopulations contribute to expansion at metastatic sites.

Overall, our findings support an integrated model of metastatic melanoma in which a subset of pre-adapted clonal “seeds,” spanning both dedifferentiated and melanocytic states, disseminate polyclonally and interact with one another and with organ-specific “soil.” Metastatic progression arises from the interplay between intrinsic lineage programs, phenotypic plasticity, and interactions within the tumor microenvironment. This framework provides a unified approach to elucidate how clonal identity and cellular state converge to drive melanoma metastatic progression.

## Methods

### Cell culture

Melanoma WND238 cell line was established in our lab, and cultured in TU2% media consisting of 80% MCDB 153, 10% Leibovitz’s L-15, 2% FBS, 2.4 mM CaCl_2_, and passaged using 0.05% trypsin-EDTA. For lentivirus packaging, we cultured Lenti-X 293T cells (Clontech) in DMEM containing 10% FBS, and similarly passaged for lentivirus production.

### Lentivirus packaging and transduction

Before plasmid transfection, Lenti-X 293T cells were grown to ∼80% confluency in 10 cm tissue culture dishes in DMEM containing 10% FBS. For each dish, the medium was replaced with 10 mL of DMEM (10% FBS) containing 25 μM chloroquine diphosphate, followed by a 5-hour incubation. For transfection, 81.6 μl of Transporter 5 (Polysciences) was added to 420 μl of Opti-MEM (Thermo Fisher Scientific, 31985062). In parallel, 8.6 μg of psPAX2, 2.6 μg of pMD2.G, and 9.2 μg of the MeRLin plasmid library were mixed in 500 μl of Opti-MEM. Diluted Transporter 5 was added dropwise to the diluted DNA tube, inverted and incubated for 20 min before being added dropwise to the culture dish^9^. Approximately 18 hours post-transfection, the medium was aspirated, cells were washed once with 1× DPBS, and fresh DMEM (5% FBS) was added. Virus-containing media were collected at 48 and 72 h post-transfection and stored at 4 °C. After the final collection, pooled virus-laden media were centrifuged at 500 × g for 5 min and filtered through a 0.45 μm PES filter. To concentrate the lentivirus, four volumes of virus-containing media were mixed with one volume of cold Lenti Concentrator (Origene, TR30025) and incubated overnight at 4 °C with constant rocking. The concentrated lentiviral particles were pelleted by centrifugation at 3,000 × g for 35 min at 4 °C and resuspended in cold, sterile 1× DPBS at 1/20th of the original media volume by gentle pipetting. The purified lentivirus was aliquoted and stored at −80 °C. To transduce melanoma cells, freshly thawed lentiviral library and polybrene (final concentration 8 μg/mL) were added to dissociated cells, which were then plated in six-well plates (225,000 cells in 1 mL of media per well) and centrifuged at 600 × g for 30 min at 32°C. After overnight incubation at 37°C, the media was removed, cells were washed once with 1× DPBS, and 2 mL of TU2% was added to each well. Cells were then passaged into 10-cm dishes in the above-mentioned media. Dissociated cells were sorted by flow cytometry for mNeptune2.5-positive cells to achieve a multiplicity of infection (MOI) below 0.4. Barcoded cells were expanded and divided into replicates, with one replicate saved as the original control.

### Animal studies

All animal procedures were approved by the Institutional Animal Care and Use Committee (IACUC, #201546). For injections of barcoded WND238 cells, 6-8-week-old male immunodeficient NSG mice were shaved on their lower back and injected subcutaneously into the right flanks (2.5 × 10^5^ in 100 μL RPMI:Matrigel (Corning), 1:1). The mice were received from the in-house breeding facility and housed under pathogen-free conditions. Tumor size monitoring was performed using caliper measurement and calculated using the following formula volume = (W2 × L)/2, where W and L refer to the short and long tumor diameter, respectively.

### Bioluminescence and fluorescence imaging

A fresh solution of D-luciferin (PerkinElmer) was prepared at 30 mg/mL in 1× DPBS and filter-sterilized through a 0.2 µm filter. Mice received an intraperitoneal injection of D-luciferin (150 mg/kg body weight) prior to imaging and were anesthetized with 2% isoflurane in oxygen. Images were acquired using the IVIS Spectrum CT system (PerkinElmer) and analyzed with Living Image 4.4 software. Dorsal and ventral scans were performed for each animal, and bioluminescence signals were captured using an open emission filter. For epi-fluorescence imaging of mNeptune2.5, excitation and emission filters of 640 nm and 680 nm were used to optimize signal-to-noise ratio. For *ex vivo* imaging, tumors and organs were immediately excised, placed in tissue culture dishes, and imaged for both bioluminescence and fluorescence signals. Tissues were cut into chunks for bulk RNA-seq, single-cell RNA-seq, and formalin-fixed paraffin-embedded (FFPE) processing.

### Immunohistochemistry

Formalin-fixed, paraffin-embedded tissue sections (1-2 μm) were stained with an anti-firefly luciferase antibody (Novus Biologicals, NB100-1677). Hematoxylin and eosin (H&E) staining was performed according to standard histological procedures.

### Bulk and single-cell sample preparation

For single-cell sample preparation, tissues were minced and added to 5 ml dissociation solution consisting 4.7 ml RPMI with 200 μl enzyme H, 100 μl enzyme R, and 25 μl enzyme A (Miltenyi Biotec, Tumor Dissociation Kit) on ice. Samples sealed in C-tubes were proceeded onto the gentleMACS Octo Dissociator (Miltenyi Biotec) with a heating apparatus for 1 h. Following dissociation, 8 ml of serum-free media was added, and single-cell suspensions were filtered through a 70 μm nylon mesh and centrifuged at 250 × g for 7 minutes. The pellet was resuspended in 1 ml ACK Lysing Buffer (Quality Biological, 118-156-101), incubated for 5 minutes, and then diluted with 9 ml serum-free media before a second centrifugation at 250 × g for 5 minutes. The pellet was resuspended in 80 µl BSA buffer (0.25 g BSA in 50 ml PBS) per 2 million cells before adding 20 μl mouse antibody cocktail per 2 million cells (Miltenyi Biotec, Mouse Cell Depletion Kit). After 15 minutes of incubation at 4°C, LS columns were primed with 3 ml BSA buffer, and the labeled cells were passed through, collecting the flow-through containing enriched human tumor cells. Cells were washed three times with 1 ml BSA buffer, centrifuged at 250 × g for 5 minutes. The supernatant was completely removed, and purified human melanoma cells were resuspended in 80 µl PBS before transferring to an Eppendorf tube on ice. A cell count was conducted before proceeding to bulk RNA-seq and scRNA-seq.

### Bulk RNA-seq

Libraries for whole transcriptome RNA sequencing were prepared using the Stranded Total RNAseq with Ribo-zero Plus kit (Illumina, San Digo, CA) as per manufacturer’s instructions starting with an input of 700 ng of total RNA and 10 cycles of final PCR amplification. Library size was assessed using the 4200 Tapestation and the High-Sensitivity DNA assay (Agilent, Santa Clara, CA). Concentration was determined using the Qubit Fluorometer 2.0 (Thermofisher, Waltham, MA). Next Generation Sequencing with a paired-end 2×150bp run length was done on the NovaSeq X platform (Illumina, San Diego, CA). A minimum of 30M reads per sample was acquired for each sample.

### Barcoded RNA-FISH

To validate the co-localization of *OLFML3* and *AXL* with the most dominant clone (barcode suffix “TCCTGCAGTA”), we used paired bulk RNA-seq and scRNA-seq to recover the full-length 265 bp barcode. A custom 5ZZ antisense probe set Syn-T1v3 (ACD 1855191-C1) was designed by the ACD Probe Design Team and synthesized to target this barcode mRNA transcript for spatial mapping. Using the RNAscopeTM Multiplex Fluorescent Reagent Kit v2 with TSA Vivid Dyes (ACD 323270), we visualized this metastatic subpopulation of barcode “TCCTGCAGTA” and the expression of *OLFML3* (RNAscope® Probe Hs-OLFML3, ACD 1124161-C2) and *AXL* (RNAscope® Probe Hs-AXL, ACD 602131-C3) in tissue sections. A 5ZZ sense probe targeting the antisense sequence of barcode “ACGACCACAA” (Syn-T2v3-sense, ACD 1587501-C1) was used as the negative control for non-specific staining^9^. These probes were used with TSA Vivid Fluorophore 650 (ACD 323273), TSA Vivid Fluorophore 570 (ACD 323272), and TSA Vivid Fluorophore 520 (ACD 323271) respectively. DAPI was used to stain nuclei. The slides were imaged using a Hamamatsu Nanozoomer S60 Whole Slide Scanner at 40× magnification. Images were stored and analyzed on the Concentriq platform (Proscia).

### Single-cell RNA-seq library preparation and read alignment

Single cell droplets were generated using the Chromium Next GEM single cell 3’ kit v3.1 (10x Genomics). cDNA synthesis and amplification, library preparation, and indexing were performed using the 10x Genomics Library Preparation kit (10x Genomics), according to the manufacturer’s instructions. The overall library size was determined using the Agilent Bioanalyzer 2100 and the high-sensitivity DNA assay, and libraries were quantitated using KAPA real-time PCR. Libraries were pooled and sequenced on the NovaSeq 6000 (Illumina, San Diego, CA, USA) using an S2 300 cycle kit (Illumina), paired end run with the following run parameters: 26 base pair × 8 base pair (dual index) × 280 base pair.

Reads alignment was performed using Cell Ranger v8.0.0 (10x Genomics). Raw sequencing reads were first aligned to a combined human-mouse reference genome (refdata-gex-GRCh38_and_GRCm39-2024-A) to identify and exclude mouse-derived reads. Cells with more than 20% of the reads mapped to the mouse genome were removed. For barcoded samples, the reads were aligned to a custom reference genome that included the barcode vector sequence integrated into the human genome (GRCh38-2024-A). All the other samples were aligned directly to the human genome (GRCh38-2024-A).

### Clonal barcode extraction

Clonal barcodes and their corresponding 10x cell barcodes (CB tags) were retrieved from the BAM files generated using Cell Ranger. Reads containing barcode sequences were aligned to a pseudo-chromosome that represented the barcode vector. We have developed a clonal barcode extraction tool to automate this process. Briefly, the extractor first scans for the known 3’ sequence of the vector backbone (CAGATCTTAGCCACTTTTTAAAAGAAAAGGGGG) to locate the start of the semi-random clonal barcode sequence in each read^9^. The extracted barcode sequences were subsequently processed through a series of quality control and correction steps including removal of UMI duplicates, empty vectors, short barcodes, barcodes with incorrect patterns, and cells containing multiple barcodes. For each high-confidence barcode, the first 10 bases and their corresponding CB tags were compiled into a structured data frame (details are available on GitHub).

### Computational analysis of scRNA-seq data

All scRNA-seq expression matrices were processed using the Seurat v5^46^. The barcode information was incorporated into the cell metadata of each Seurat object. Cells were filtered out if they had fewer than 2,000 detected genes, more than 50,000 unique molecular identifiers (UMIs), or over 20% of the reads mapped to mitochondrial genes. Doublets were identified and removed using DoubletFinder v2.0.4^47^. To assess the clonal dynamics of barcoded cells over time, we calculated the Shannon diversity index^11^ for each sample based on the barcode abundance distributions. Cell cycle phase scores were calculated based on the canonical S and G2/M genes. After quality control, the samples were integrated and normalized using SCTransform^48^.

To better resolve the clonal architecture within each scRNA-seq sample, we used ClonoCluster v0.0.1^13^, an algorithm that integrates both transcriptomic profiles and clonal barcode identities into hybrid clusters. This approach replaced standard Louvain clustering and uniform manifold approximation and projection (UMAP) in the Seurat workflow, which did not produce optimal cluster separation due to the absence of clonal barcode information in its analysis. To fine-tune the impact of clonal barcode identity on the UMAP layout, we tested various Warp Factor (WF) values and determined that WF = 6 provided the best balance between transcriptomic similarity and clonal architecture.

### Functional signature analysis by barcode group

To characterize the transcriptional programs and functional signatures associated with distinct clonal populations, we redefined the ClonoCluster-derived clusters into barcode groups. Each group comprised a unique and mutually exclusive set of barcodes.

For each barcode group, we performed differential gene expression analysis to identify upregulated genes (fold change ≥ 1.5, detected in ≥ 40% of cells, and adjusted p-value < 0.05). These genes were subsequently submitted to the Enrichr platform for functional enrichment analysis^49^. Hallmark, KEGG, Reactome, and Wikipathways databases were queried to determine biological processes associated with each group’s transcriptional program.

After defining the representative functional signatures for each group, we selected key genes from the enriched pathways and calculated signature activity scores using AUCell^50^ to validate the activity of these group-specific programs at single-cell level. Cell-cell communication between groups was inferred using CellChat v2.1.2^51,52^.

### SCENIC analysis

To infer the transcription factor activity and gene regulatory networks for each barcode group in the scRNA-seq data, we performed single-cell regulatory network inference and clustering (SCENIC) analysis using pySCENIC v0.12.1^50^. Gene co-expression modules were first inferred using the GRNBoost2 algorithm^53^. These modules were then refined by filtering target genes based on motif enrichment within ±10 kb of transcription start sites using the hg38 motif collection as a reference. Next, regulon activity was quantified across all cells using AUCell for each regulon. The resulting activity scores were integrated into Seurat objects to identify the top differentially active regulons across barcode groups defined from ClonoCluster.

### Alternative polyadenylation analysis

To investigate the post-transcriptional regulation associated with treatment response, we performed alternative polyadenylation analysis (APA) using MAAPER^54^ by comparing BAM files from drug treated samples to non-treated samples (day 21, day 57, endpoint vs. day 0; BRAFi/MEKi vs. CTRL). The required polyadenylation site (PAS) annotation file for hg38 was obtained from https://github.com/Vivianstats/data-pkg/tree/main/MAAPER/PolyA_DB. Genes with an adjusted p-value < 0.05 and log fold change > log (1.2) or < −log (1.2) were classified as lengthened or shortened, respectively.

### Computational analysis of bulk RNA-seq data

Raw sequencing data were aligned to the human reference genome (hg38) using STAR v2.7.11b^55^ and gene-level counts were quantified. Transcript abundance was normalized as transcript per million (TPM) for single-sample gene set enrichment analysis (ssGSEA)^56^. The enrichment scores of the functional signatures identified in each barcode group were evaluated.

### Statistical tests

All statistical analyses were performed using R v4.4.0. Only post hoc false discovery rate (FDR) adjusted p-values less than 0.05 were considered significant.

## Data availability

Raw and processed scRNA-seq and bulk RNA-seq data were deposited in Gene Expression Omnibus (GEO) database and will be made publicly available upon peer reviewed publication. Reviewer access will be provided upon request.

## Code availability

The analysis and visualization scripts for this study are available at GitHub (https://github.com/Yeqing95/MeRLin), along with a brief guide on reproducing the analysis workflows and figures presented in the paper.

## Additional Information

Supplementary Information is available for this paper.

## Acknowledgements

The authors thank Wistar Genomics Facility, especially S. Majumdar and S. Widura, for assistance with bulk and single-cell RNA sequencing; Wistar Histotechnology Facility, especially F. Chen and X. Wang, for assistance with paraffin-embedding and staining; Wistar Microscopy Facility, especially F. Keeney, for assistance with microscopy; Wistar Animal Facility, especially D. DiFrancesco, M. Houston-Leslie, B. Grant, and E. Conicello, for assistance with mouse experiments; Wistar Flow Cytometry Facility, especially J. Faust and J. Fundyga, for assistance with flow sorting; G. Robertson at Penn State Cancer Institute, A. L. Welm at University of Utah, D. Yu at MD Anderson Cancer Center, for discussions on the project. This research was supported by NIH/NCI grants P01 CA114046-15, R01 CA238237-05, R01 CA240362-05, R01 CA241148-05, P50 CA261608-03, U54 CA224070-05, and the Sheldon G. Adelson Medical Research Grant R-202206-00540.

## Author contributions

H.L. and M.H. conceived and designed the study. H.L. designed and performed all experiments and acquired, analyzed, and interpreted the data under the supervision of M. H.. Y.C. analyzed all data and contributed to data interpretation. M.D., N.P., and H.L. contributed to the production and transduction of lentivirus. M.D., N.P., J.K., H.L., M.X., and Q.Z. supported cell cultures. M.D. and H.L. conducted the flow cytometry. H.L., J.K., N.P., D.F., M.D., G.S.B., M.X., and M.T. performed mouse experiments. H.L., J.K., and M.D. prepared the bulk, single-cell, and FFPE samples. M.D., and L.L. performed microscopy. Y.C. conducted the bulk and single-cell RNA-seq data analyses, SCENIC analysis and APA analysis. C. Q. analyzed clonal frequency. X.X. provided guidance on melanoma metastasis models. N.A., A.R., Z.W., X.X., D.S.H., B.T., and J.V. provided feedback on the early manuscript drafts. H.L. wrote the manuscript with input from all the authors.

## Declaration of interests

The authors declare no competing interests.

## Notes

### Competing Interest Statement

The authors have declared no competing interest.

