## Supplementary Figures for "Single-cell lineage tracing maps clonal and transcriptional dynamics in melanoma metastasis"

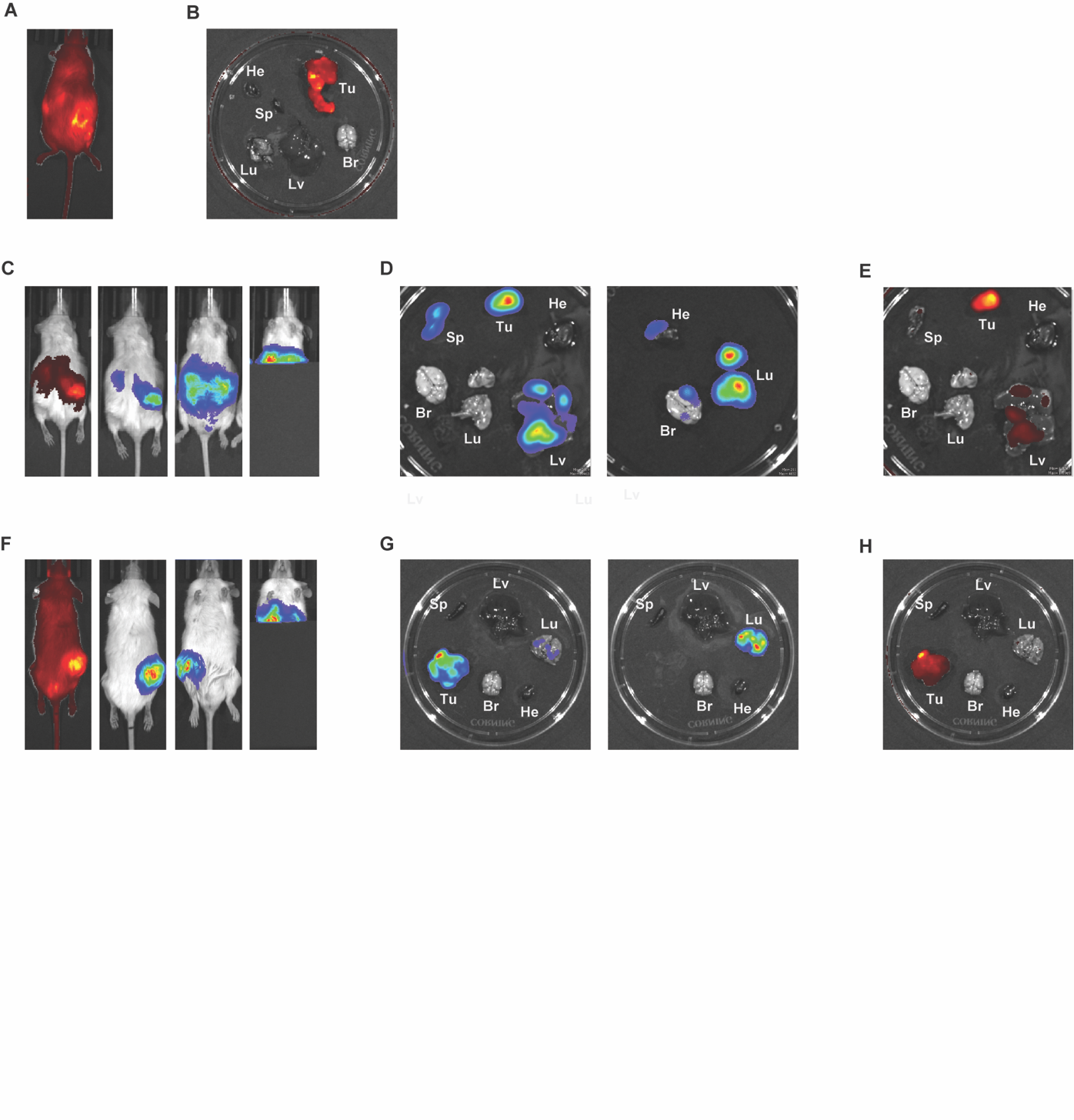
**Fig. S1** *In vivo* and *ex vivo* imaging of metastatic dissemination in MeRLin-barcoded WND238 melanoma. **A**, *In vivo* fluorescence imaging of the first mouse bearing a barcoded WND238 tumor. **B**, *Ex vivo* fluorescence imaging of the excised primary tumor (Tu) and organs, including lung (Lu), liver (Lv), spleen (Sp), brain (Br), and heart (He) of the first mouse. **C**, *In vivo* fluorescence (left) and bioluminescence imaging (BLI, second to right) of the second mouse in dorsal (left and second) and ventral (third and right) views. After shielding the primary tumor with a light proof cover, residual signals are detected in the lungs (right). **D**, Ex vivo BLI of the excised primary tumor and organs of the second mouse, imaged with (left) and without (right) the primary tumor, liver, and spleen. **E**, Corresponding *ex vivo* fluorescence imaging of (**D** left). **F**, *In vivo* fluorescence (left) and BLI (second to right) of the third mouse in dorsal (left and second) and ventral (third and right) views. Shielding the primary tumor revealed residual signals in the lungs (right). **G**, Ex vivo BLI of the excised primary tumor and organs of the third mouse, imaged with (left) and without (right) the primary tumor. **H**, Corresponding *ex vivo* fluorescence imaging of (**G** left).

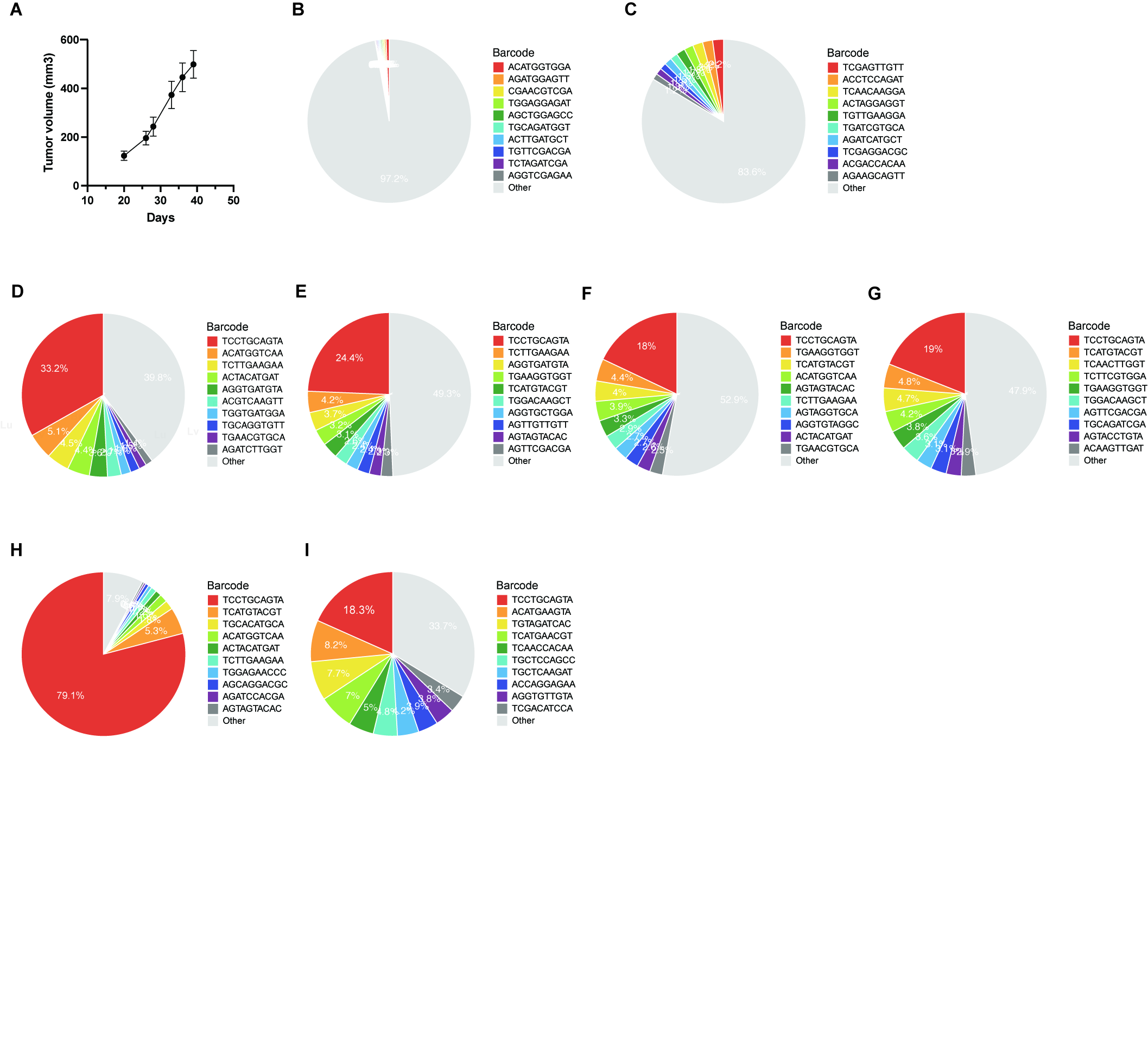

**Fig. S2** Clonal composition in metastatic and non-metastatic melanoma models. **A**, In vivo growth curves of barcoded WM4237-1 tumors. **B**, Pie chart showing the clonal composition of the original control of barcoded WM4237-1 cells. **C**, Pie chart showing the clonal composition of WM4237-1 tumor following engraftment. **D-G**, Pie charts showing the hierarchical clonal composition of the primary tumor (**D**), lung metastases (**E**), liver metastases (**F**), and spleen metastases (**G**) from the second mouse (Table S3, bulk RNA seq). **H-I**, Pie charts showing the hierarchical clonal composition of the primary tumor (**H**) and lung metastases (**I**) from the third mouse (Table S3, bulk RNA seq).

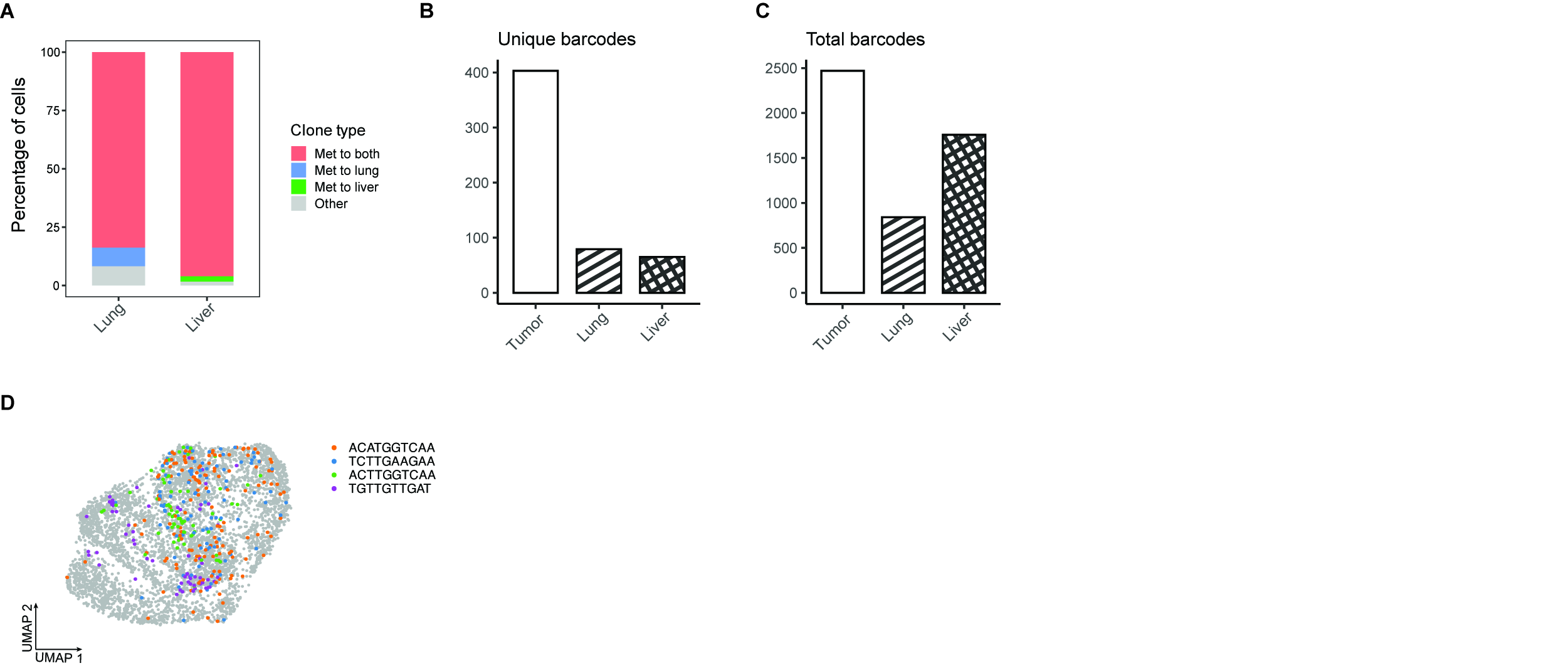

**Fig. S3** Clonal composition and metastatic fate across primary and metastatic sites. **A**, Cell percentage of metastatic fate categories in lung and liver metastases. **B**, Numbers of unique barcodes detected in the primary tumor, lung, and liver metastases. **C**, Total barcode counts detected in the primary tumor, lung, and liver metastases. **D**, UMAP visualization of the metastatic fates of subpopulations ranked 2 to 5 within the primary tumor.

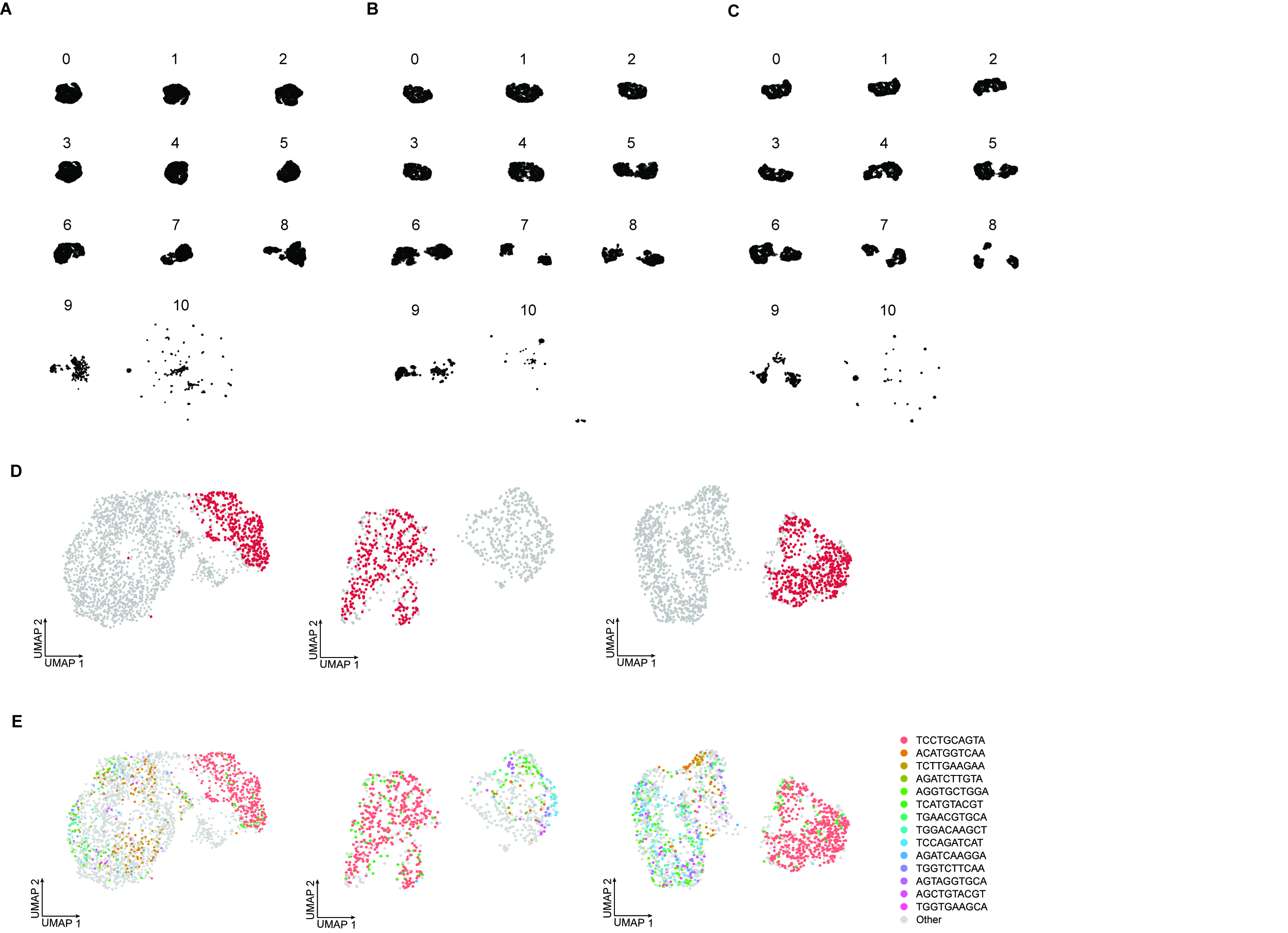

**Fig. S4** ClonoCluster integration of clonal and transcriptomic information. **A-C**, Effect of increasing Warp Factor values on UMAP structure for scRNA-seq data from the primary tumor (**A**), lung metastases (**B**), and liver metastases (**C**), illustrating the progressive contribution of clonal identity to hybrid clustering. **D**, UMAPs showing the distribution of cells carrying the most abundant barcode in the primary tumor (left), lung metastases (middle), and liver metastases (right). **E**, UMAPs showing the distribution of cells carrying the 14 shared barcodes in the primary tumor (left), lung metastases (middle), and liver metastases (right).

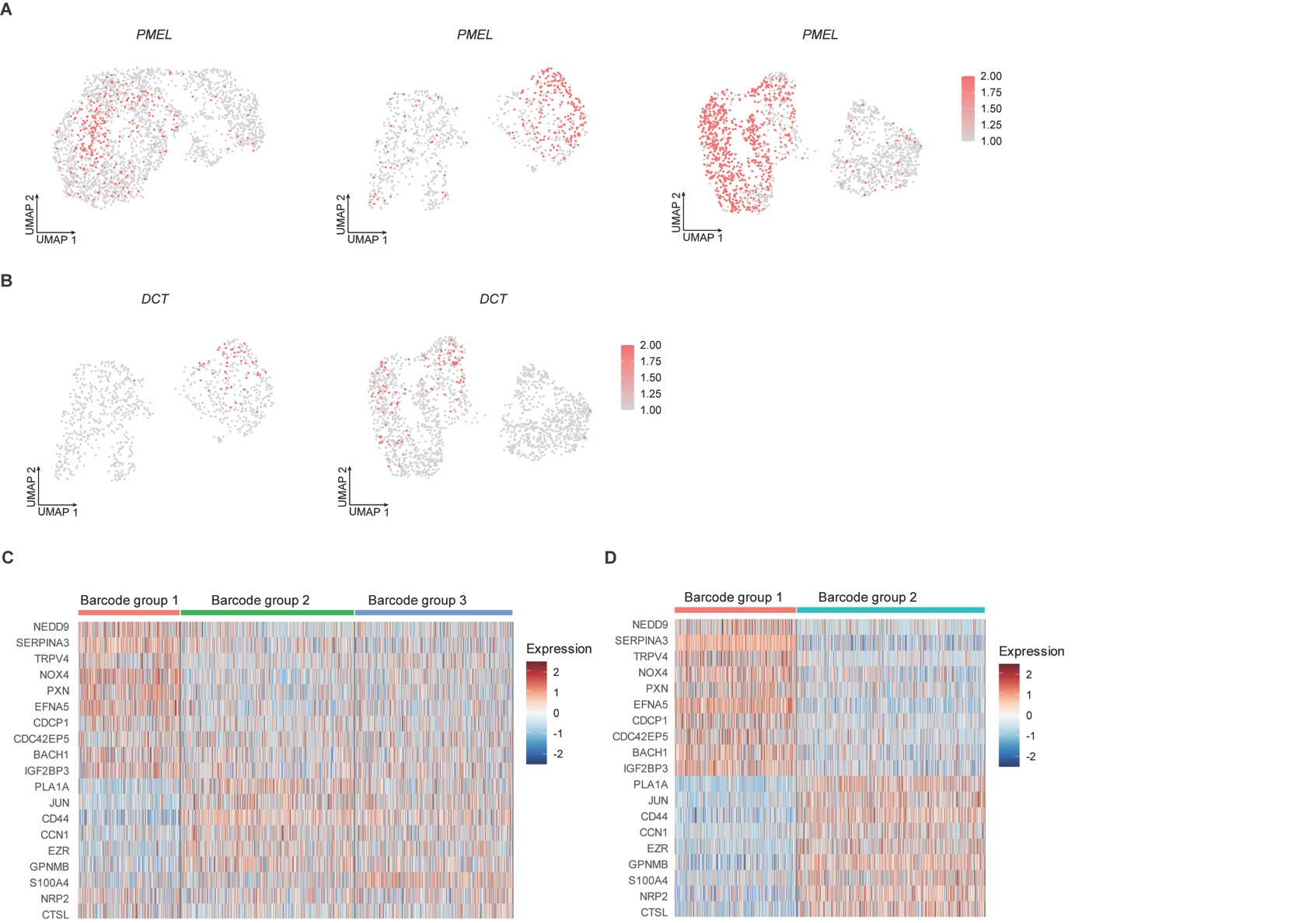

**Fig. S5** Melanocytic marker and invasion-associated gene expression across primary and metastatic melanoma. **A**, UMAPs showing the melanocytic marker *PMEL* expression in the primary tumor (left), lung metastases (middle), and liver metastases (right). **B**, UMAPs of the melanocytic marker *DCT* in lung metastases (left) and liver metastases (right). **C**, Heatmap showing melanoma invasion-associated genes identified for each barcode group in the primary tumor and liver metastases (**D**) (Table S4). Fold change ≥ 1.5, cell percentage ≥ 40%, and adjusted P < 0.05.

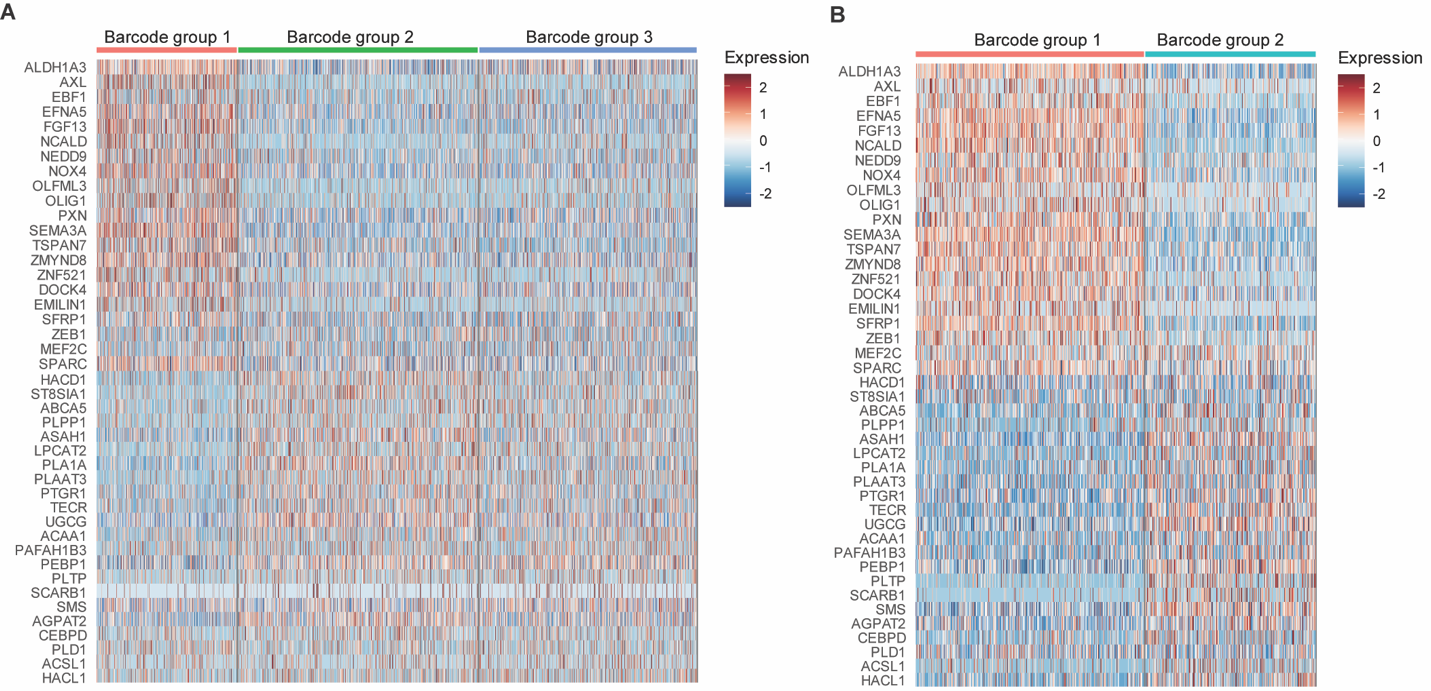

**Fig. S6** NC-like and lipid metabolism signatures across barcode groups in primary tumor and lung metastases. **A**, Heatmap showing neural crest stem-like (NC-like) and lipid metabolism signature genes identified for each barcode group in the primary tumor and lung metastases (**B**) (Table S5). Fold change ≥ 1.5, cell percentage ≥ 40%, and adjusted P < 0.05.

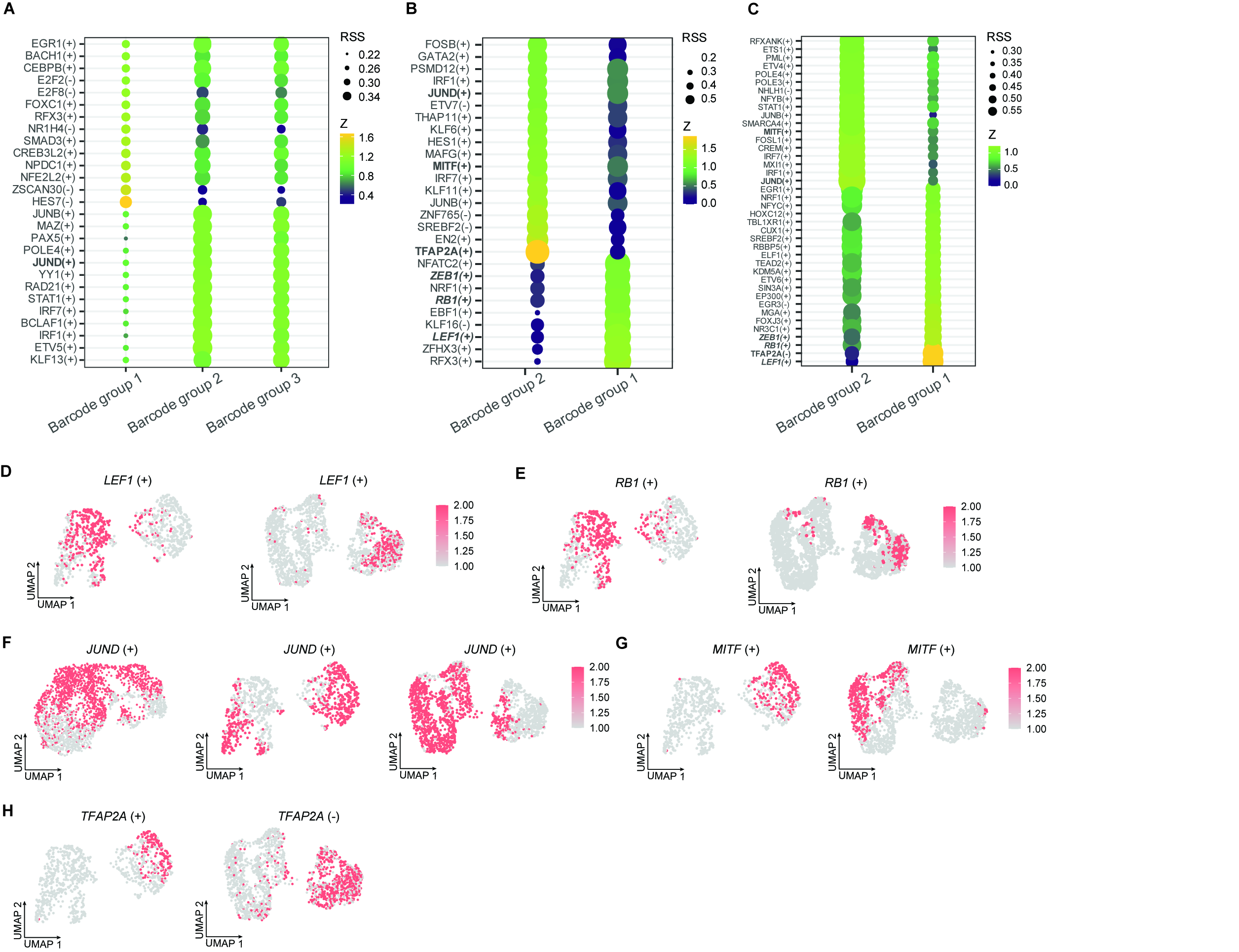

**Fig. S7** Distinct regulon activities in barcode groups across primary and metastatic sites. **A,** SCENIC analysis identifying key regulators of each barcode group in primary tumor, lung metastases (**B**), and liver metastases (**C**). **D**, SCENIC analysis showing regulon activities of transcription factors *LEF1* and *RB1* (**E**) in lung metastases (left) and liver metastases (right). **F**, Regulon activities of transcription factor *JUND* across primary tumor (left), lung metastases (middle), and liver metastases (right). **G**, Regulon activities of transcription factors *MITF* and *TFAP2A* (**H**) in lung metastases (left) and liver metastases (right).

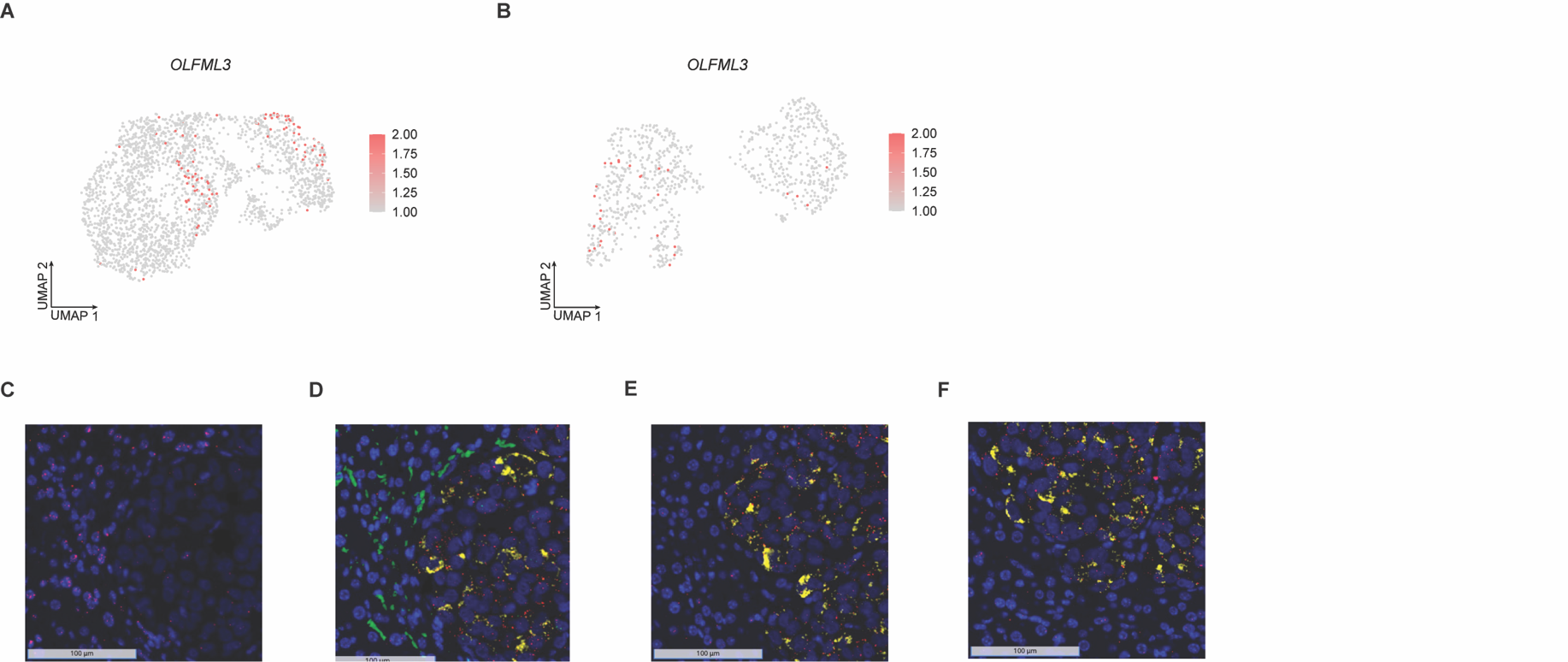

**Fig. S8** *In situ* localization of a dominant metastatic subpopulation and *OLFML3* expression. **A**, UMAPs of the NC-like marker *OLFML3* in primary tumor and lung metastases (**B**). **C**, RNA-FISH showing the staining of non-specific negative control barcode probes (red) in liver tissue (scale bar 100 μm). DAPI staining was used to visualize cell nuclei (blue). **D**, RNA-FISH showing merged view of a dominant metastatic subpopulation (red, barcode suffix “TCCTGCAGTA”), human *OLFML3* expression (yellow) and staining of probes targeting human *AXL* (green) in liver tissue (scale bar 100 μm). **E-F**, ROIs showing merged partial co-localization of the dominant metastatic subpopulation (red, barcode suffix “TCCTGCAGTA”) and *OLFML3* expression (yellow) at tumor-liver interface (scale bar 100 μm).
